# Microstructural Mapping of Neural Pathways in Alzheimer’s Disease using Macrostructure-Informed Normative Tractometry

**DOI:** 10.1101/2024.04.25.591183

**Authors:** Yixue Feng, Bramsh Q. Chandio, Julio E. Villalon-Reina, Sophia I. Thomopoulos, Talia M. Nir, Sebastian Benavidez, Emily Laltoo, Tamoghna Chattopadhyay, Himanshu Joshi, Ganesan Venkatasubramanian, John P. John, Neda Jahanshad, Robert I. Reid, Clifford R. Jack, Michael W. Weiner, Paul M. Thompson, Alzheimers Disease Neuroimaging Initiative

## Abstract

**Introduction:** Diffusion MRI is sensitive to the microstructural properties of brain tissues, and shows great promise in detecting the effects of degenerative diseases. However, many approaches analyze single measures averaged over regions of interest, without considering the underlying fiber geometry.

**Methods:** Here, we propose a novel Macrostructure-Informed Normative Tractometry (MINT) framework, to investigate how white matter microstructure and macrostructure are jointly altered in mild cognitive impairment (MCI) and dementia. We compare MINT-derived metrics with univariate metrics from diffusion tensor imaging (DTI), to examine how fiber geometry may impact interpretation of microstructure.

**Results:** In two multi-site cohorts from North America and India, we find consistent patterns of microstructural and macrostructural anomalies implicated in MCI and dementia; we also rank diffusion metrics’ sensitivity to dementia.

**Discussion:** We show that MINT, by jointly modeling tract shape and microstructure, has potential to disentangle and better interpret the effects of degenerative disease on the brain’s neural pathways.

## 1 INTRODUCTION

Diffusion MRI (dMRI) is a powerful approach often used to investigate the microstructure and geometry of the brain’s neural pathways; it measures the characteristics of water diffusion in the living brain ^1,2^. As water diffusion is constrained along the brain’s neural pathways,fiber tracking (or tractography) using diffusion profiles detected with dMRI can reconstruct the 3D geometry of the brain’s major fiber bundles. Pathological processes that occur in degenerative diseases, such as the loss of neurons and myelin and inflammation, affect tissue diffusion properties to alter both tissue microstructure and pathway geometry. Because of this, dMRI is sensitive to pathological processes that are not detectable with standard anatomical MRI.

Anisotropy and diffusivity measures computed from diffusion tensor imaging (DTI) are the most widely used measures to characterize white matter (WM) microstructural properties. These diffusion metrics have been studied in degenerative, developmental, and psychiatric conditions ^3^. As many types of molecular pathology influence the dMRI signal, including amyloid and tau protein accumulation in the brain, a large literature has focused on charting WM anomalies that arise in the progression of neurodegenerative disorders such as Alzheimer’s disease (AD) ^4–6^, Parkinson’s disease ^7^ and other dementias. Thomopoulos *et al*. ^5^ examined four standard DTI metrics and how they related to dementia severity in 730 individuals scanned as part of the Alzheimer’s Disease Neuroimaging Initiative (ADNI). They found that mean diffusivity (MD) was associated with age and dementia severity, assessed using the clinical dementia rating (CDR) scale. A follow-up study ^6^ examined dMRI metrics in the cortical gray matter. They found that cortical dMRI metrics mediated the relationship between cerebrospinal fluid (CSF) markers of AD and delayed logical memory performance, which is typically impaired in early AD. Lower CSF A*β*142 and higher pTau181 were associated with cortical dMRI measures reflecting less restricted diffusion and greater diffusivity; this apparent link between AD pathology and diffusion metrics has bolstered interest in using dMRI to study AD. Even so, standard analysis methods typically reduce microstructural indices to summaries over relatively large regions of interest. These limitations have stimulated efforts to map disease effects on brain microstructure along WM tracts at a finer anatomical scale ^8,9^.

DTI-based metrics such as fractional anisotropy (FA), radial diffusivity (RD), and axial diffusivity (AxD) are susceptible to fiber crossings ^10^ —their values are influenced by the presence of multiple fiber populations in a single voxel ^11^ and not specific to any individual fiber population. While voxel-, fixel-based ^12^ and tractometry approaches ^*‡*^ have been proposed to further improve inter-subject alignment and help resolve crossing fibers, many microstructural measures are still calculated at the voxel level. Additionally, current tractometry methods often compute group statistics on for each bundle separately using a univariate approach, without accounting for the complex pattern of intersecting fibers in the brain. Tractography data can also be used to study macrostructural, or ‘shape’ properties of WM bundles. Schilling *et al*. ^13^ computed bundle shape metrics in a large multi-site cohort of elderly adults and found heterogeneous patterns of age-related macrostructural changes in the brain WM compared to the more homogeneous patterns of microstructural changes. A recent study ^14^ found that early tau-related WM changes in AD are macroscopic using fixel-based analysis metrics. To our knowledge, no work has examined how WM micro- and macro-structure are jointly altered in neurodegenerative conditions such as AD using tractometry methods, which we address in the current study.

In this study, we propose *Macrostructure-Informed Normative Tractometry (MINT)* to jointly model microstructural measures and co-occurring variations in fiber bundle geometry, with a deep learning method called the variational autoencoder (VAE). When used as a normative model, VAE can encode the anatomical patterns of normal variability for diffusion metrics in healthy controls. This multivariate model integrates multiple complementary microstructural features and accounts for the statistical covariance among different dMRI metrics as well as along-tract spatial correlations. We compare MINT-derived measures with traditional univariate measures derived from DTI, to investigate characteristic patterns of WM abnormalities in mild cognitive impairment (MCI) and dementia in a large multi-site sample. We also examine how WM abnormalities relate to clinical measures of dementia severity. As there is interest in identifying the best microstructural metrics for detecting and tracking dementia, we also rank DTI metrics by evaluating their sensitivity to dementia. After visualizing WM microstructural abnormalities in dementia and MCI in two cohorts of different ancestry and demographics, we examine how they relate to overall tract geometry, noting ambiguities of interpretation that can be resolved by joint statistical modeling of microstructure and shape.

## 2 MATERIALS AND METHODS

### 2.1 Participant Demographics

In total, we analyzed diffusion data of 1032 subjects from two cohorts —730 participants from the Alzheimers Disease Neuroimaging Initiative (ADNI) collected using 7 acquisition protocols, and an independent sample of 302 Indian-ancestry participants, evaluated and scanned at two scanners at the National Institute of Mental Health and Neuro Sciences (NIMHANS) in Bangalore, India. The subset of the ADNI cohort that was scanned with dMRI consisted of 447 cognitively normal participants (CN), 214 diagnosed with mild cognitive impairment (MCI) and 69 with dementia. For the purposes of the current study, we analyzed their baseline scans only, although most of these participants also have longitudinal dMRI data. The NIMHANS cohort consists of dMRI data from 123 CN participants, 89 people with MCI, and 90 with dementia. The demographics information for all subjects categorized by protocols and scanners are detailed in in Table 1.

**TABLE 1.** Demographics for the ADNI and NIMHANS data used in this study.

| PROTOCOL | TOTAL N | N(CN) | N(MCI) | N(DEMENTIA) | AGE MEAN ± STD | MALE (%) |
| --- | --- | --- | --- | --- | --- | --- |
| ADNI S31 | 96 | 59 | 27 | 10 | 72.03 ± 8.51 | 39 (40.62%) |
| ADNI S55 | 270 | 175 | 71 | 24 | 74.41 ± 7.93 | 120 (44.44%) |
| ADNI S127 | 112 | 68 | 36 | 8 | 73.66 ± 7.58 | 53 (47.32%) |
| ADNI P33 | 56 | 37 | 14 | 5 | 75.74 ± 7.68 | 34 (60.71%) |
| ADNI P36 | 46 | 20 | 23 | 3 | 73.14 ± 6.93 | 24 (52.17%) |
| ADNI GE36 | 42 | 21 | 17 | 4 | 72.24 ± 7.15 | 24 (57.14%) |
| ADNI GE54 | 108 | 67 | 26 | 15 | 76.11 ± 8.06 | 55 (50.93%) |
| ADNI Total | 730 | 447 | 214 | 69 | 74.13 ± 7.93 | 349 (47.81%) |
| NIMHANS Siemens | 192 | 91 | 45 | 56 | 67.02 ± 8.26 | 111 (57.81%) |
| NIMHANS Philips | 110 | 32 | 44 | 34 | 67.65 ± 7.13 | 66 (60.0%) |
| NIMHANS Total | 302 | 123 | 89 | 90 | 67.25 ± 7.86 | 177 (58.61%) |
| Total | 1032 | 570 | 303 | 159 | 72.12 ± 8.5 | 526 (50.97%) |

### 2.2 Data Preprocessing

#### 2.2.1 Alzheimer‘s Disease Neuroimaging Initiative (ADNI)

One of the two datasets analyzed in this article was obtained from the ADNI database (adni.loni.usc.edu). The ADNI was launched in 2003 as a public-private partnership, led by Principal Investigator, Michael W. Weiner, MD. The primary goal of ADNI has been to test whether serial magnetic resonance imaging (MRI), positron emission tomography (PET), other biological markers, and clinical and neuropsychological assessments can be combined to measure the progression of mild cognitive impairment (MCI) and early Alzheimer‘s disease (AD).

We analyzed dMRI data from phase 3 of ADNI, which recently concluded, from 730 participants, collected using 7 acquisition protocols - S31, S55, S127, P33, P36, GE36 and GE54 (Table 1) ^*§*^. The acquisition protocols and preprocessing pipelines are detailed in ^5,17^.

#### 2.2.2 National Institute of Mental Health and Neuro Sciences (NIMHANS)

For the first scanner used at NIMHANS —a Philips 3T Ingenia scanner—dMRI was acquired using a single-shot, diffusion-weighted echo-planar imaging sequence (TR=7441 ms, TE=85 ms, TA (total acquisition time) =630 s, voxel size: 2*×*2*×*2mm^3^, 64 slices, flip angle=90^*°*^, FOV 224mm). Detailed descriptions of the acquisitions and quality assurance can be found in Parekh *et al*. ^18^. For the Siemens 3T Skyra scanner, dMRI data was also collected using a singleshot, diffusion-weighted echoplanar imaging sequence (TR=8400 ms, TE=91 ms, TA (total acquisition time) = 546 s, FOV 240mm) ^19,20^. For both scanners, transverse sections of 2-mm thickness were acquired parallel to the anterior commissure-posterior commissure (AC-PC) line, and diffusion weighting was encoded along 64 independent spherical directions, using a *b*-value of 1000 s/mm^2^. Preprocessing steps for all DWI volumes in the NIMHANS cohort closely followed the ADNI preprocessing pipeline ^5^, and included: denoising using local principal component analysis ^21^ implemented in DiPy ^22^, Gibbs ringing removal ^23,24^ implemented in DiPy ^22^, eddy currents ^25^ using FSL’s *eddy_cuda* ^26^ and bias field inhomogeneity ^27^ using MRtrix’s *dwibiascorrect*. The NIMHANS Philips data was acquired with reverse phase-encoded blips (b0s only), and FSL’s *topup* was used to estimate the susceptibility-induced distortion ^28^. As the NIMHANS Siemens protocol did not acquire dMRI with blip-up blip-down acquisitions, Synb0-DisCo ^29^ was used to generate synthetic non-distorted b0 images, to estimate the distortions with *topup*. After processing dMRI, diffusion tensors were fitted at each voxel, using the non-linear least-squares method to produce FA, MD, RD, and AxD scalar maps in the dMRI native space. Fiber orientations were reconstructed using Robust and Unbiased Model-BAsed Spherical Deconvolution (RUMBA-SD) along with Contextual Enhancement ^30,31^. Particle filtering tracking ^32^ was used to generate whole-brain tractograms, with 8 seeds per voxel generated from the WM mask, a step size of 0.2 mm, angular threshold of 30^*°*^, and the continuous map stopping criterion ^32^.

#### 2.2.3 TractoInferno

As explained below, we used a deep learning method to encode the mathematical properties of normal statistical variation in the brain’s WM bundles, assisted by auxiliary information from another high quality dataset. In the terminology of deep learning, such an auxiliary dataset is referred to as a pre-training dataset, which can help robustly encode and estimate normal variation in DTI measures and tract geometry in healthy subjects. In our study, this pre-training dataset consisted of 198 single-shell dMRI volumes already preprocessed from the publicly available TractoInferno training dataset ^33^, acquired on 3T scanners from 6 sites. Constrained spherical deconvolution ^34^ and deterministic tracking ^22^ were used to generate whole-brain tractograms. For all ADNI, NIMHANS and the TractoInferno datasets, DiPy’s ^22^ auto-calibrated RecoBundles method ^8,16^ was used to segment 30 WM bundles using the population-averaged HCP-842 atlas ^15^ in both the native and MNI (Montreal Neurological Institute) space, see Figure 1.

**FIGURE 1.**
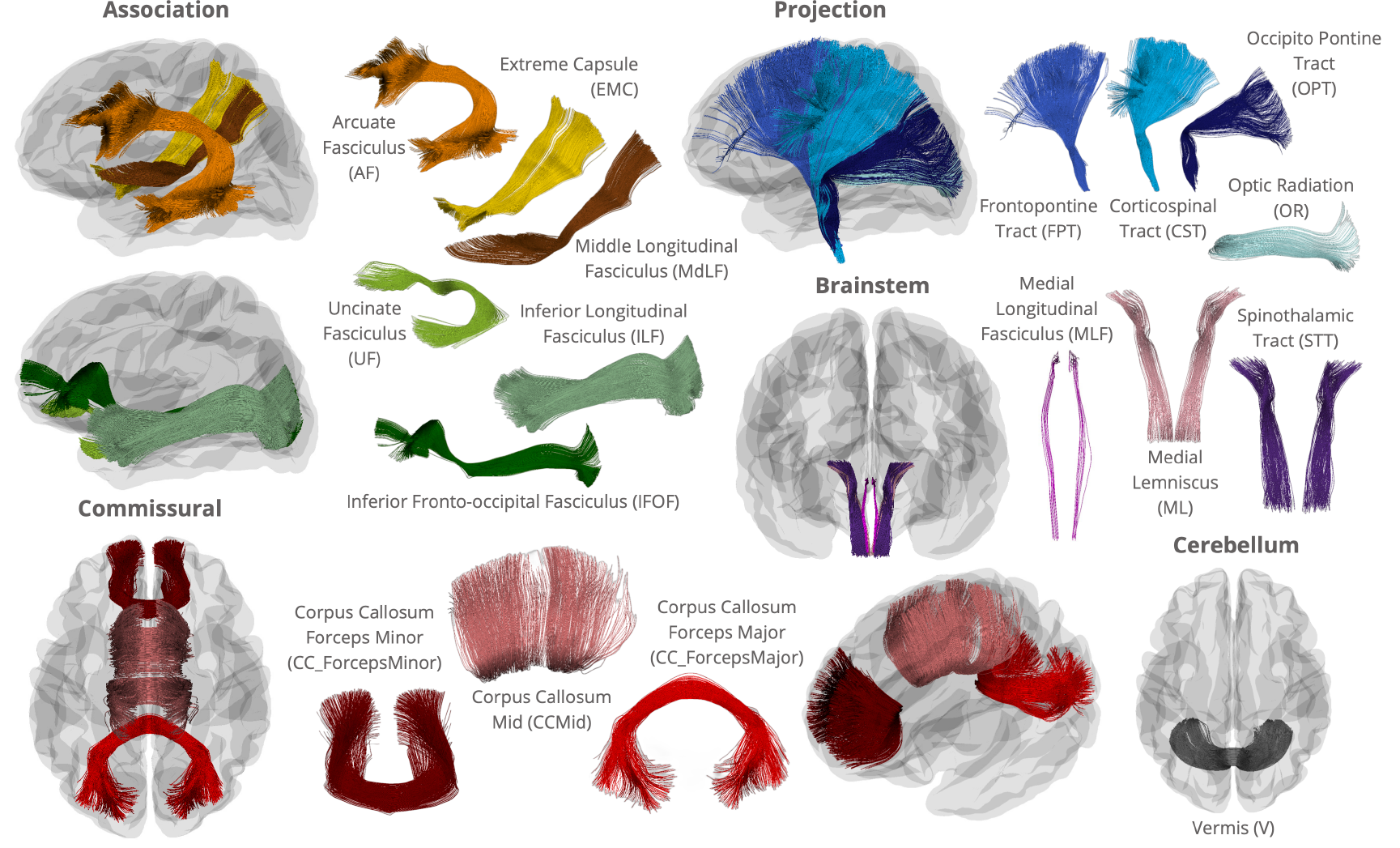
WM bundles from the HCP842 atlas ^15^ used in RecoBundles ^16^ in this study. Bundles visualized individually outside of the glass brain are not to scale. While bilateral association and projection bundles are used in this study, only those in the left hemisphere are shown.

### 2.3 Macrostructure-Informed Normative Tractometry (MINT)

#### 2.3.1 Macrostructure Modeling

In our Macrostructural-Informed Normative Tractometry (MINT) framework, a key component is the variational autoencoder (VAE) model, which learns patterns of macrostructural (tract shape and geometry) variation in a healthy population based on the 3D streamline coordinates provided by tractography in addition to the microstructural features along those coordinates. VAE ^37^ is considered to be one of many nonlinear extensions of dimensionality reduction methods, such as principal components analyses (PCA), where the variation in a multidimensional dataset is often encoded using a small number of latent factors. These underlying factors represent a probability distribution over the input data and enable statistical inference in a fully generative model.

In MINT, we use 1D convolutional layers to model streamline-level information. This method was proposed to enforce sequential dependency of 3D points along a streamline ^38^. Illustrated in Figure 2(a), a 1D convolution learns from local neighbor-hood points along a streamline using a ‘sliding window’, and multiple convolutional layers can be stacked to learn information from points farther away. This approach has been successfully applied to problems such as streamline filtering, clustering ^39^ and generative sampling of streamlines for bundle segmentation ^40^.

**FIGURE 2.**
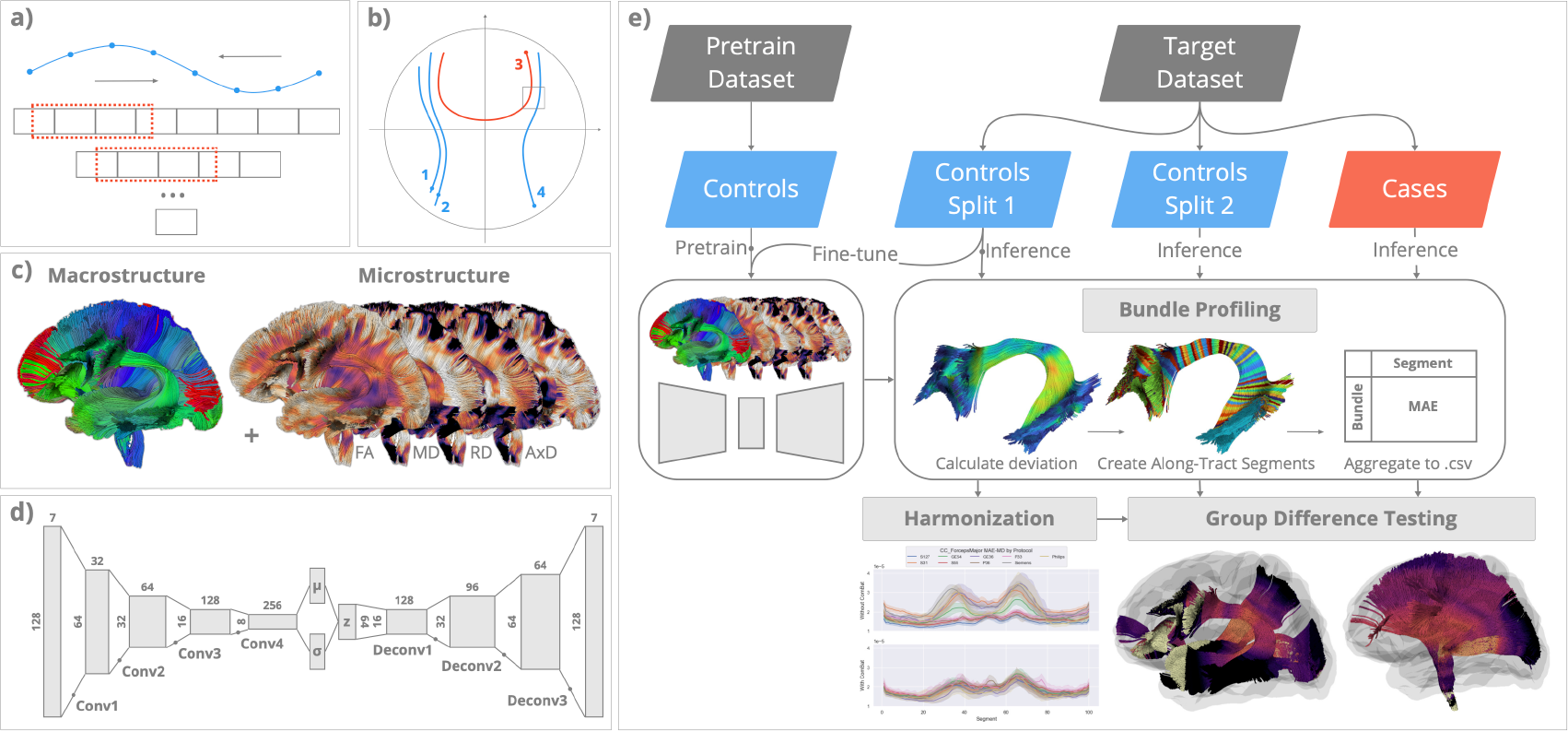
Modeling whole-brain macro- and micro-structure in MINT. **(a)** MINT uses 1D convolutional layers in a variational autoencoder (VAE) to model the sequential dependency along each streamline; **(b)** MINT models whole-brain fiber geometry by training the VAE on streamlines from multiple fiber bundles. Proximity and similarity between streamlines are encoded using global coordinates; **(c)** Whole-brain macrostructure is represented using tractography data, and whole-brain microstructure is represented by projecting scalar maps of DTI metrics onto every point on each streamline. Both types of data are combined to train the VAE model. **(d)** The VAE is a generative model, trained via backpropagation, to reconstruct its input data as accurately as possible while passing all the intermediate data through a narrow bottleneck layer (*z* shown in the middle). VAE enforces the data to be optimally compressed into a latent set of parameters, and the means *μ* and covariance *σ* of these parameters follow a multivariate Gaussian distribution. This enables the rich complexity of variations in fiber microstructure and 3D geometry to be modeled using a compact representation. **(e) The MINT framework**. A variational autoencoder (VAE) model is first trained on healthy controls from a pretraining dataset (TractoInferno). For each target dataset (ADNI, NIMHANS), the controls are split into 2 sets (Split 1 and Split 2) and Split 1 is used to fine-tune the VAE model. We then perform model inference on all data from the target dataset to obtain their reconstruction for subsequent bundle profiling. For each bundle, the point-wise deviation score (MAE) is indexed using 100 along-tract segments created to align data across subjects. MAE for all points within each segment is averaged to create a bundle profile of deviation scores. In the harmonization step, bundle profiles created from data in Split 1 are used to train ComBat ^35,36^, whose parameters are applied to bundle profiles from healthy controls in Split 2 and the cases (participants with MCI and AD). Harmonized bundle profiles are then used for group difference testing using linear regression.

MINT can learn the patterns of statistical variation in whole-brain macrostructure by training it on streamlines (3D curves) from multiple bundles. While the prior literature has used the term *macrostructure* to describe bundle-level shape information, here we use this term to describe the overall, whole-brain fiber geometry derived from tractography (including information from multiple bundles). We illustrate this in Figure 2(b) showing examples of projection fibers (1,2,4) and commissural fibers (3). If each streamline was modeled independently, with rotational invariance enforced, 1D convolution would be able to distinguish streamline 1,2,4 from streamline 3, due to their different lengths and curvatures. Still, it may have difficulty distinguishing between streamlines 2 and 4. On the other hand, if streamlines from multiple bundles are normalized with respect to the whole brain, the model can learn that streamlines 1 and 2 are more likely to belong to the same bundle and share similar microstructural properties than streamlines 1 and 4, because of their global coordinates. In the case of a fiber crossing, illustrated here for streamlines 3 and 4, the model can encode information on their spatial proximity. Through joint statistical modeling of micro- and macro-structure, variance introduced by the underlying fiber geometry - that may also influence microstructural measures such as FA - could be attributed to a difference in macrostructure instead. We previously showed that a VAE with 1D convolutional layers (ConvVAE) preserves streamline and bundle distances in the latent space by training on only 3D coordinates using this approach ^41^. Through unsupervised training, the model can learn that streamlines with similar shapes and in similar positions likely belong to the same bundle.

As shown in Figure 2(c), MINT is trained on whole-brain macrostructure as well as the 4 standard DTI microstructural measures. By applying whole-brain normalization, MINT leverages the flexibility offered by streamline-based deep neural networks during training and inference. The model can produce a reconstruction for any collection of streamlines during inference, and the deviation score derived from the reconstruction error is calculated for every point on each streamline. This tactic of using reconstruction error from a VAE, or a related generative model, has been widely employed for anomaly detection in both biomedical and industrial contexts ^42,43^. Next, we create along-tract segments aligned across subjects to localize tract-specific deviations. The methods to create along-tract profiles for these deviations are detailed in Section 2.3.3.

#### 2.3.2 Data Preparation and Training

Streamline coordinates used for training were derived from 30 bundles extracted in the MNI (Montreal Neurological Institute) space. DTI metrics, FA, MD, RD, and AxD, are projected onto the corresponding bundle in the native space. Each streamline is resampled to 128 equidistant points —so that the resulting set of points can be used with 1D convolutional layers —with 7 input features - *x, y, z*, FA, MD, RD and AxD. To preserve the spatial relationships between streamlines within each subject, unit sphere normalization was applied to the 3D coordinates for all streamlines in each subject’s bundles, using a centroid and radius calculated from the HCP-842 atlas tractogram ^15^. Min-max scaling was applied to each of the four DTI metrics using preset ranges, and all 7 input features were scaled between -1 and 1. As streamlines are also ‘flip-independent’ —that is, one streamline can be traversed in either direction —we performed random flipping as an augmentation step ^40^ to further enforce this property, although 1D convolution can learn information from neighboring points in both directions.

As the VAE model is used here in a normative model setting, we need to be careful not to over-parameterize the model to avoid reconstructing anomalies at inference time. 1D depthwise separable convolution (SepConv) ^44^ was used in the encoder and decoder to accommodate a deeper feature space with efficient use of model parameters. Overall, the VAE model consists of 464,408 parameters: the encoder consists of four 1D SepConv (separable convolution) layers with kernel sizes of 15, and 32, 64, 128 and 256 channels; the decoder consists of three 1D SepConv layers with the same kernel sizes of 15, and 128, 96 and 64 channels (see Figure 2(d)). This asymmetric architecture, where the decoder is smaller than the encoder, was adopted to encourage the model to learn more robust representations ^45^ while reducing the number of parameters. The model was trained with the ELBO (evidence-based lower bound) loss ^37^. We used cyclical annealing on the KL (Kullback-Leibler) divergence term ^46^ to prevent posterior collapse, a common issue during VAE training. The pretraining data, consisting of streamlines from 30 bundles from 198 subjects in the TractoInferno dataset, contains over 100 million streamlines. To improve computational efficiency during data loading, the VAE model sampled a batch of 512 streamlines, with replacement, for each parameter update, and was pretrained for 24,000 steps.

For both the ADNI and NIMHANS datasets, data from the cognitively normal (CN) participants were split into independent subsets of size 35%/15%/50% for training/validation/testing. Data from participants with MCI or dementia were concatenated with the 50% CN subjects to create the test set. The test set contained a large percentage of healthy control subjects, to better estimate group difference statistics from the resulting deviation scores. The validation set was used to adjust the model’s fine-tuning parameters, such as batch size and number of training steps, and inspect the reconstruction quality. The pretrained VAE model was fine-tuned on 30 bundles from the subjects in the training set, in each cohort, for 5000 steps. We did not pool the training data in ADNI and NIMHANS together in this step, to simulate the scenario where sites cannot share raw data, but can still share bundle profiles computed from their own fine-tuned model. A complete diagram of the MINT framework —including training, inference, bundle profiling, and group difference testing —is shown in Figure 2(e).

#### 2.3.3 Bundle Profiling

At inference time (i.e., when the model is applied to new datasets without updating its parameters), each bundle *X* from the test set of participants in both ADNI and NIMHANS is passed through its respective fine-tuned model, to obtain their reconstruction *X*^*′*^ of the same shape. We then use the assignment map approach from BUAN ^8^ to create 100 along-tract segments, aligned across subjects. Every point in *X* is assigned a segment index (1 to 100), and the same index is applied to the reconstructed bundle. We then calculate the deviation using Mean Absolute Error (MAE) for each segment *i* and feature *k* as follows:

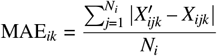

Here *N*_*i*_ is the number of points assigned to segment *i* in *X*, and the features include shape (calculated from 3D coordinates), FA, MD, RD and AxD. To compare the MAE scores with the original DTI microstructural measures, we also calculate the mean along-tract profile *M* as follows

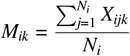

where the features contain 4 DTI measures. For each subject’s bundle, *X*, we created along-tract profiles for a total of 9 features: MAE scores were calculated for 5 features at 100 along-tract segments, which we will refer to as MAE-Shape, MAE-FA, MAE-MD, MAE-RD and MAE-AxD, to distinguish them from the mean along-tract DTI profile, calculated for 4 feature at the same 100 along-tract segments - DTI-FA, DTI-MD, DTI-RD and DTI-AxD. The same bundle profiling procedures were repeated for the training set in both cohorts, for ComBat harmonization (detailed next).

Along-tract bundle profiles for all 9 features were harmonized using ComBat using the bundle-wise approach described in Chandio *et al*. ^17^. ComBat is a ‘location and scale’ model, where we assume that each measurement is affected by the additive and multiplicative effects of the scanner and protocol ^36^, estimated using an empirical Bayes method. Covariates can also be specified in ComBat to preserve the known biological variability across subjects (e.g. age and sex). In this study, ComBat was run for each of the 30 bundles and 9 features, by pooling information across 100 segments of each bundle, with age and sex modeled as covariates. We used the Python implementation of ComBat (https://github.com/Jfortin1/neuroCombat). To prevent data leakage ^47,48^, we estimated ComBat parameters on bundle profiles created from the training set for both cohorts previously used in fine-tuning. The training set contained only CN subjects, so the diagnosis was not included in the ComBat model during training. The estimated ComBat parameters were then applied to bundle profiles from the test set, which included CN and subjects with MCI and dementia, for subsequent statistical analysis.

### 2.4 Statistical Analysis of Diagnostic Group Differences

To evaluate effects of MCI and dementia on MINT-derived along-tract MAE and raw DTI measures and how they differ between the ADNI and NIMHANS cohorts, we performed statistical analyses on the harmonized bundle profiles of all 9 features for both the dementia and MCI groups, compared to the relevant group of control participants, using linear regression separately for each cohort. A linear regression model was fitted for each of the 100 segments in 30 bundles with diagnosis (DX), age, and sex modeled as independent variables. We corrected for multiple comparisons using the false discovery rate (FDR) method ^49^ applied to 3000 tests across the whole brain. Only bundle profiles from the test set were used for statistical analysis in this step and Section 2.5.

### 2.5 Associations with Clinical Metrics

To assess how MAE and raw DTI measures are associated with standardized clinical measures in the ADNI cohort, we computed partial correlations between a standard dementia severity rating, the CDRsb (clinical dementia rating, sum-of-boxes score), and 9 bundle profile features, removing the effect of age, and sex. The CDRsb measures are widely used as a primary outcome measure in clinical trials of anti-amyloid treatments and other interventions ^50^. Each bundle profile from the ADNI test set was averaged to obtain one scalar value per bundle per feature, and participants with a missing CDRsb score were removed. Correlation was calculated using the Spearman rank-order correlation, and we report *ρ* correlation coefficient and its 95% confidence interval. We used the partial correlation implementation from the Pingouin Python package, version 0.5.4^51^.

## 3 RESULTS

### 3.1 Visualizing MCI and Dementia Effects on White Matter Tracts

In Figures 3, 4, 5, we visualize along-tract group differences (i.e., the regression coefficient for the effect of diagnosis on microstructure, *β*(DX) of DTI and MAE measures), for both MCI vs. CN and dementia vs. CN comparisons in the ADNI and NIMHANS cohorts ^*¶*^. All visualizations in these figures were created using the FURY package ^22,52,53^. As expected from the intensification and spread of disease pathology between MCI and dementia, we see greater effects of dementia than MCI in almost all WM regions, for all 9 measures. Overall, mean diffusivity (MD), which is commonly reported as increased in dementia, was higher in MCI and higher still in dementia. Consistent with the temporal lobe atrophy that is typical of both MCI and dementia, the abnormal elevation in mean diffusivity showed strongest effects in the temporal lobe pathways (ILF, temporal UF and AF bundles), and in the corpus callosum at the midline.

**FIGURE 3.**
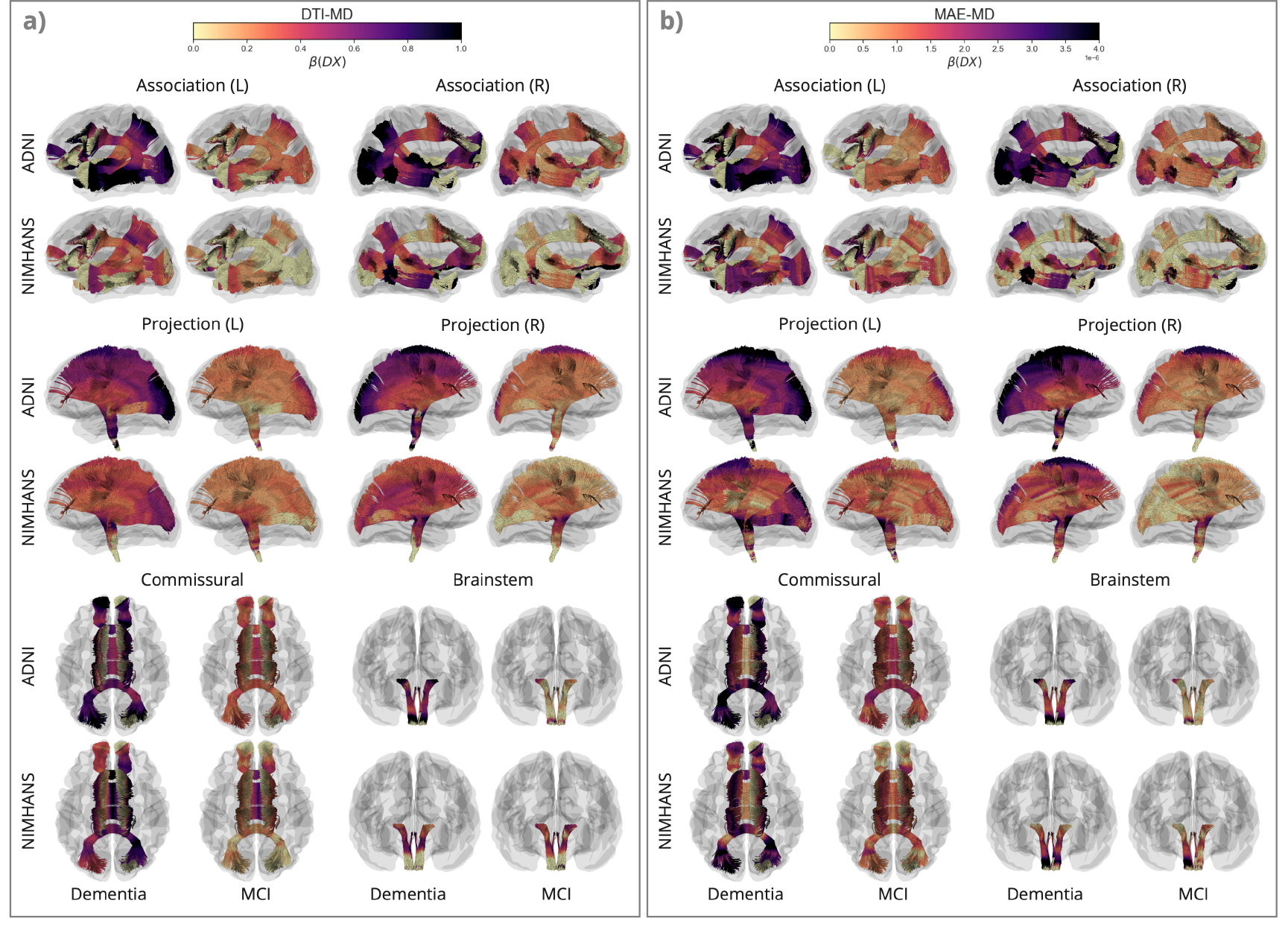
Effects of Dementia and Mild Cognitive Impairment (MCI) on Mean Diffusivity (MD). Here, we show along-tract effects of diagnostic group, by color-coding the regression coefficients associated with diagnostic group, *β*(DX), for DTI-MD and MAE-MD, for dementia vs. CN and MCI vs. CN in the ADNI and NIMHANS cohorts, categorized by WM pathways. The colors are mapped using different scales for MAE and raw DTI values, as MAE measures - calculated from reconstruction error - generally are smaller in magnitude than the DTI measures. The plots show patterns of effects but are not intended for direct comparison of effect sizes across measures, except across 3 corresponding diffusivity measures where the color map range is the same.

**FIGURE 4.**
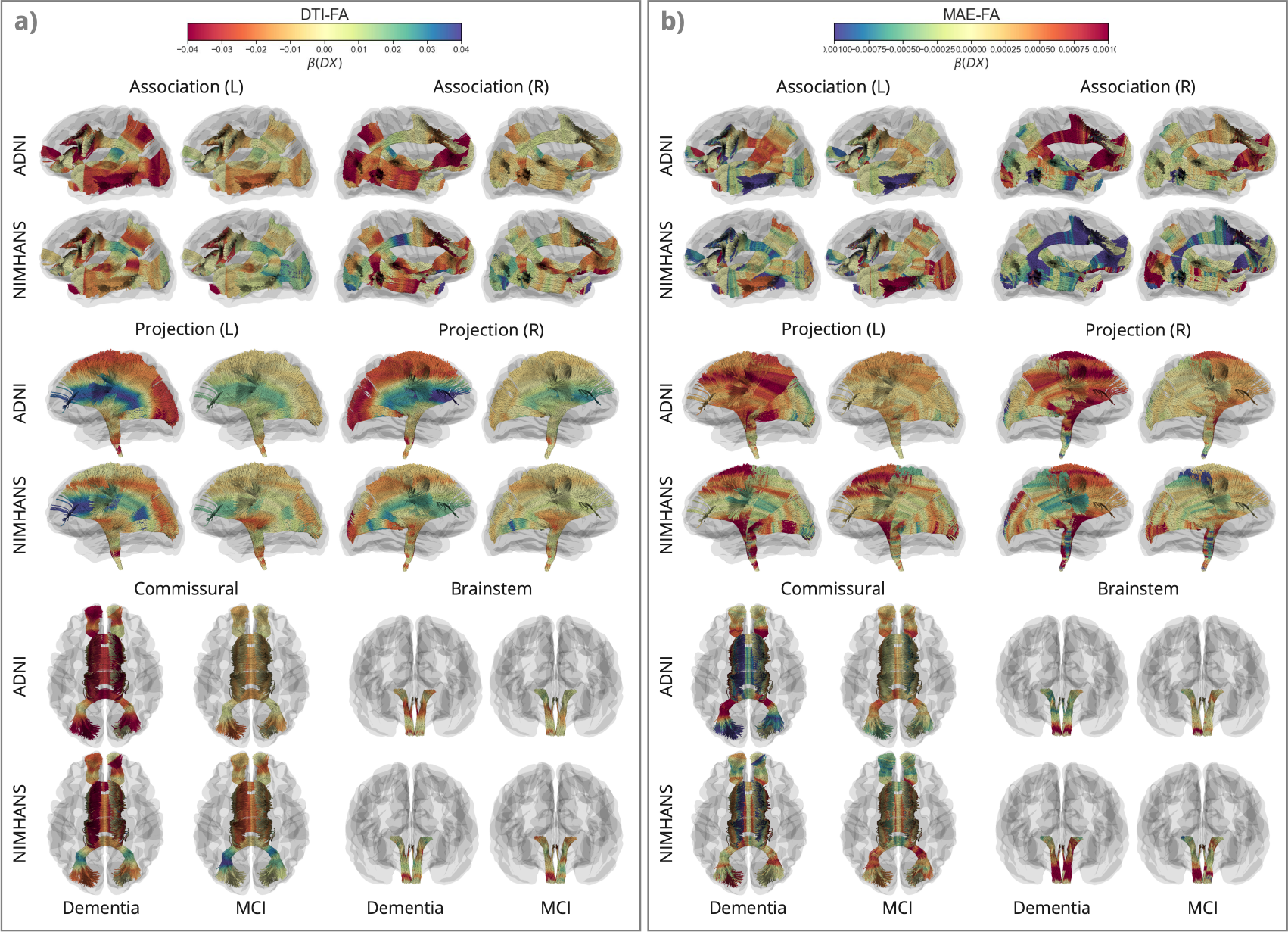
Effects of Dementia and Mild Cognitive Impairment (MCI) on Fractional Anisotropy (FA). Along-tract *β*(DX) for DTI-FA and MAE-FA, for dementia vs. CN and MCI vs. CN in the ADNI and NIMHANS cohorts, categorized by WM pathways. The color map is flipped for DTI-FA compared to MAE-FA and MAE-Shape in Figure 5, to better illustrate disease effects on these measures (*in red*).

**FIGURE 5.**
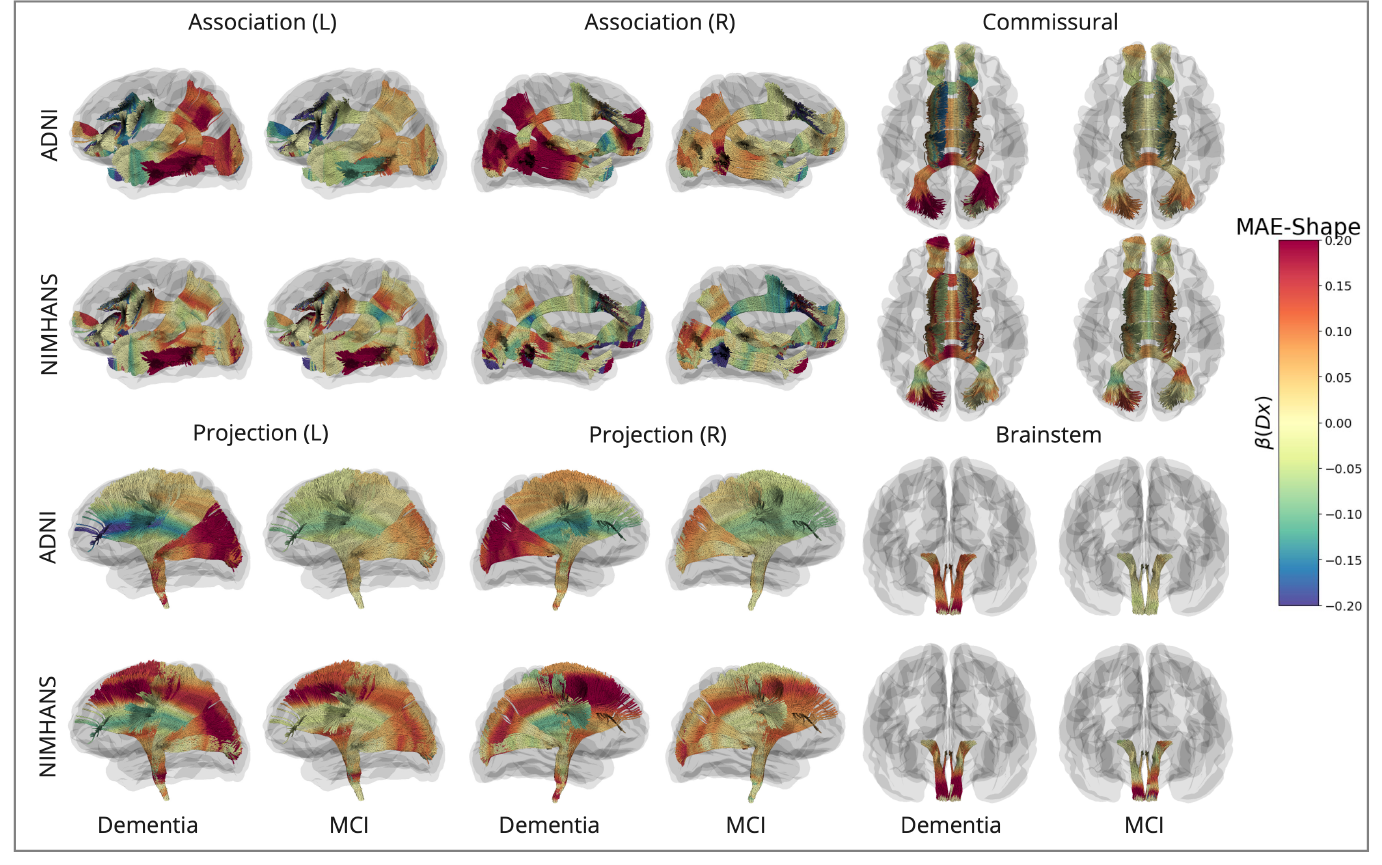
Effects of Dementia and Mild Cognitive Impairment (MCI) on MAE-Shape. Along-tract *β*(DX) for MAE-Shape, for dementia vs. CN and MCI vs. CN in the ADNI and NIMHANS cohorts, categorized by WM pathways.

For diffusivity measures in general, higher DTI-MD, DTI-RD, DTI-AxD, MAE-MD, MAE-RD, and MAE-AxD were found in dementia and MCI compared to control participants. Lower DTI-FA, higher MAE-FA, and higher MAE-Shape (denoting a bundle’s geometrical distortion compared to its normal configuration learned from the VAE model) are found in dementia and MCI compared to CN, in most brain regions. For all MAE metrics, these findings are consistent with our expectation that statistical deviation from the norm is greater in the MCI and dementia groups than in the CN group. Trends and directions of deviation for DTI metrics are also consistent with prior reports on regional DTI measures in AD ^54,55^. Patterns of dementia and MCI effects were also mostly consistent across the two cohorts we examined, with stronger statistical significance in ADNI compared to NIMHANS.

For MAE-MD and DTI-MD (Figure 3), we see widespread effects of dementia across the whole brain, with the largest effects in the posterior and temporal regions, including the OPT and OR bundles in the projection pathways, the ILF, posterior regions of the MdLF and EMC and temporal regions of the UF and AF bundles in the association pathways, and the *forceps major* in the commissural pathways. The patterns are largely similar for both DTI-MD and MAE-MD, and across the two cohorts. We see smaller effects in NIMHANS than ADNI, except for the brainstem bundles —where effects could be inflated due to the ComBat harmonization (**Supplementary Section 5**).

Without considering crossing fibers, the overall diffusivity can be decomposed into a diffusivity along the axons (AxD), and a component transverse to axons (RD). In the context of neurodegeneration, the latter is increased when cell membranes are damaged, neurons are lost, or when myelin degrades. MCI and dementia effects on MAE-RD were similar to those found for MAE-MD in all regions (**Supplementary Figure S1(b)**), whereas their effects on MAE-AxD and MAE-MD are more similar in some bundles in the association, brainstem and projection pathways (**Supplementary Figure S2(b)**). The ADNI cohort showed greater MCI and dementia effects with stronger significance for the three MAE diffusivity measures in most regions relative to the NIMHANS cohort. The difference across cohorts is most pronounced for MAE-AxD. Compared to ADNI, in the NIMHANS cohort, we found a smaller effect of dementia on MAE-AxD in the temporal association pathways; this cross-cohort difference is larger compared to MAE-MD and MAE-RD measures in the same regions. In the commissural pathways, however, the NIMHANS cohort shows a higher effect of dementia on DTI-AxD compared to ADNI in the *forceps major* and the middle section of the CCMid bundle.

For the univariate DTI measures, all three diffusivity measures were increased in both dementia and MCI across the 2 cohorts, with similar patterns in the association, commissural, and brainstem pathways across measures. Projection pathways showed a widespread elevation in DTI-MD in both dementia and MCI, but the patterns of increased DTI-RD and DTI-AxD differ along the lengths of the tracts. The sharp increase of RD from the proximal parts closer to the deep nuclei and brainstem to the distal parts closer to the cortex(**Supplementary Figure S1(a)**) —and the increase of DTI-AxD in the opposite direction (**Supplementary Figure S2(a)**) —result in a fan-shaped pattern of the projection pathways. This pattern is most pronounced for DTI-FA (see Figure 4(a)), where FA is lower in the distal parts in dementia, and higher in the proximal parts for both dementia and MCI.

This ‘fan-shaped’ pattern is more prominent for the raw DTI metrics (FA, RD, and AxD) than their counterpart MAE metrics. The primary difference between raw and MAE metrics for the DTI measures is that they are modeled jointly along with macrostructure in MINT. Interestingly, patterns of dementia effects on DTI-FA are more similar to dementia effects on MAE-Shape, see Figure 5 than on MAE-FA. The ‘fan-shaped’ pattern in the projection pathways is seen for both MAE-Shape and DTI-FA, where the proximal parts show lower MAE-Shape and higher DTI-FA, and distal parts show higher MAE-Shape and lower DTI-FA in both dementia and MCI. Aside from the projection pathways, the similarity between MAE-Shape and DTI-FA is also seen in the posterior and temporal regions of the association pathways and the *forceps major* in the commissural pathways. After accounting for the effects of whole-brain macrostructure using MINT, patterns of dementia and MCI differences in MAE-FA show the greatest deviation from their raw DTI counterpart, compared to the three diffusivity measures. Most bundle segments that showed significant group differences in DTI-FA no longer pass the significance threshold after FDR correction for MAE-FA; the remaining significant differences are primarily in the projection and commissural pathways (**Supplementary Figure S6**).

The ‘fan-shaped’ pattern in the projection pathways is seen in both ADNI and NIMHANS cohorts for DTI-RD, DTI-AxD, and DTI-FA, with highly similar patterns across cohorts. For MAE measures, however, MCI and dementia participants from the ADNI cohort show a more homogeneous increase in MAE-RD, MAE-AxD, and MAE-FA along-tract, whereas the along-tract patterns are not consistent in the NIMHANS cohort. Interestingly, the NIMHANS cohort shows larger dementia and MCI effects on MAE-Shape in the distal parts of the projection pathways compared to ADNI, but smaller effects in the association and commissural pathways.

### 3.2 Association of Diffusion Metrics with Dementia Severity

We plot the partial Spearman rank-order correlation coefficient *ρ* and its 95% confidence interval for each bundle, to assess the association between nine diffusion derived measures (described in Section 2.3.3) averaged over 100 segments per bundle and CDRsb measures of dementia severity, in Figure 6. We compare the four DTI-derived measures with their corresponding MINT-derived measures, as well as MAE-Shape. Overall, both the raw DTI and MAE diffusivity measures are largely similar, where each metric shows a significant positive correlation with CDRsb in most bundles, except IFOF_L and V. MAE diffusivity measures align the most with raw DTI measures for MD, followed by RD and AxD, where the association with RD measures is slightly higher for MAE in the association and projection pathways, and the association with AxD measures is slightly lower for MAE in the projection pathways. For FA measures, there is a weak correlation between MAE-FA and CDRsb, lower than that for DTI-FA in most bundles, except EMC and FPT, where they are comparable or higher. MAE-Shape shows both positive and negative correlation with CDRsb, where all association and brainstem bundles exhibit positive correlations except for UF, ML_R, and V. The partial correlation for MAE-Shape varies more in the projection and commissural bundles, consistent with findings in Section 3.1, where dementia and MCI effects on MAE-Shape vary along these bundles and are less prominent in MAE-FA compared to DTI-FA.

**FIGURE 6.**
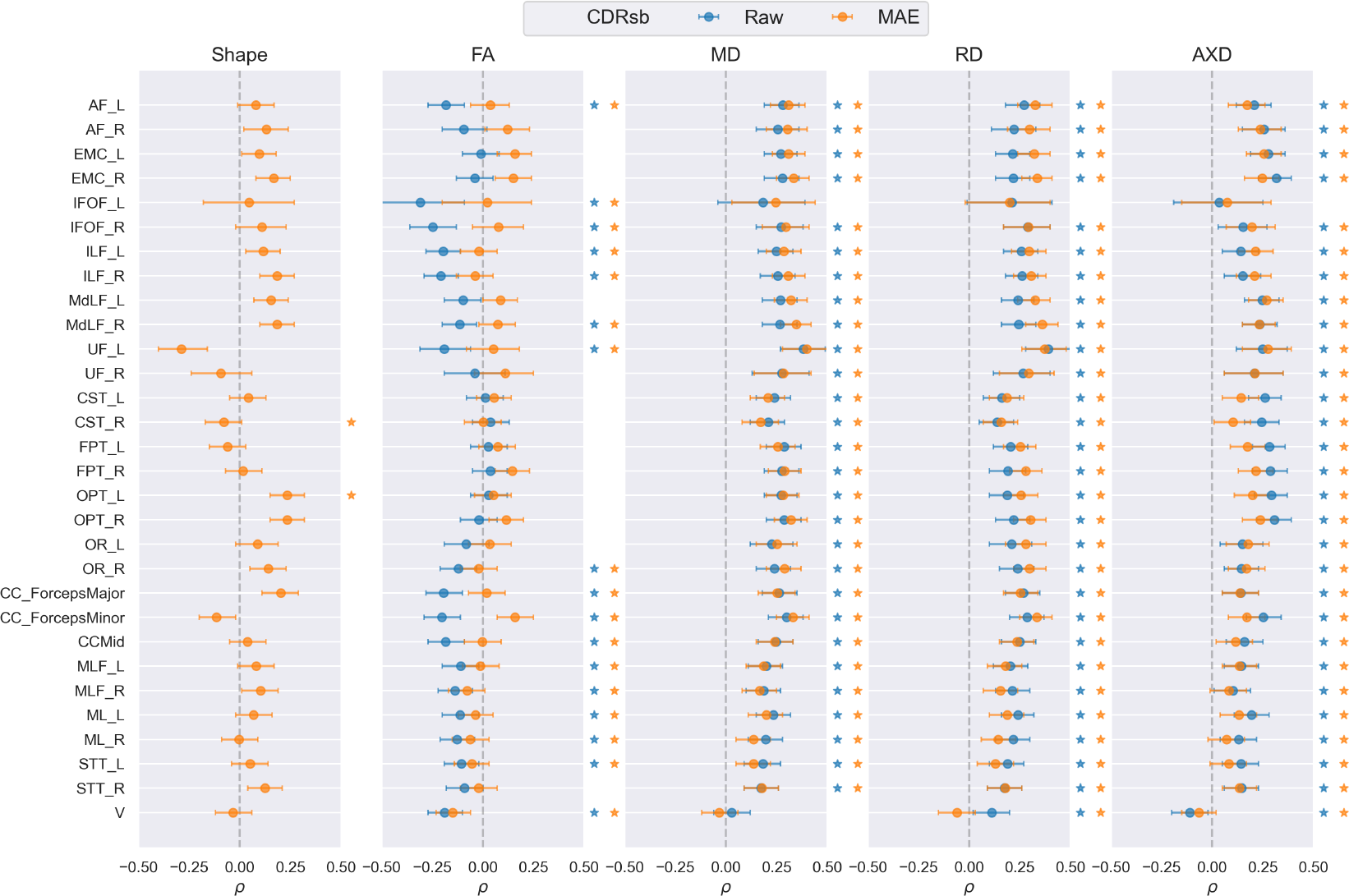
Partial Spearman rank-order correlation coefficient (*ρ*) and its 95% confidence interval for all 30 bundles between CDRSB score and 9 diffusion measures, calculated in Section 2.3.3. The corresponding raw and MAE measures for each DTI metric are shown in the same panel and MAE-Shape is shown separately. Correlation coefficients that are statistically significant after FDR correction are marked with a star to the right of each plot with the corresponding color.

### 3.3 Comparing DTI- and MINT-Derived Diffusion Measures

There is considerable interest in determining which dMRI-derived metrics are most strongly associated with dementia. In other words, if we could rank the metrics by their strength of association with dementia, we could decide which ones are the most effective biomarkers in picking up subtle and distributed effects of the disease in the brain. As the statistics are defined at multiple points along a tract, the standard way to compare effect sizes in this scenario is to use a Quantile-Quantile (QQ) plot to gauge how much of the tract shows significance and at what level of significance. To evaluate how DTI- and MINT-derived measures differ in detecting group differences, we show the QQ of – log_10_(*P*) for the dementia vs. CN comparison in the ADNI cohort in Figure 7. These plots are cumulative histograms of the *P* values of association between dementia diagnosis and each of the diffusion metrics. They can be used to rank the strength of association for different brain biomarkers, and summarize both the strength and extent of significant associations. If a curve rises sharply from the origin in the QQ plot, it means that a large proportion of the tract has very low *P* values.

**FIGURE 7.**
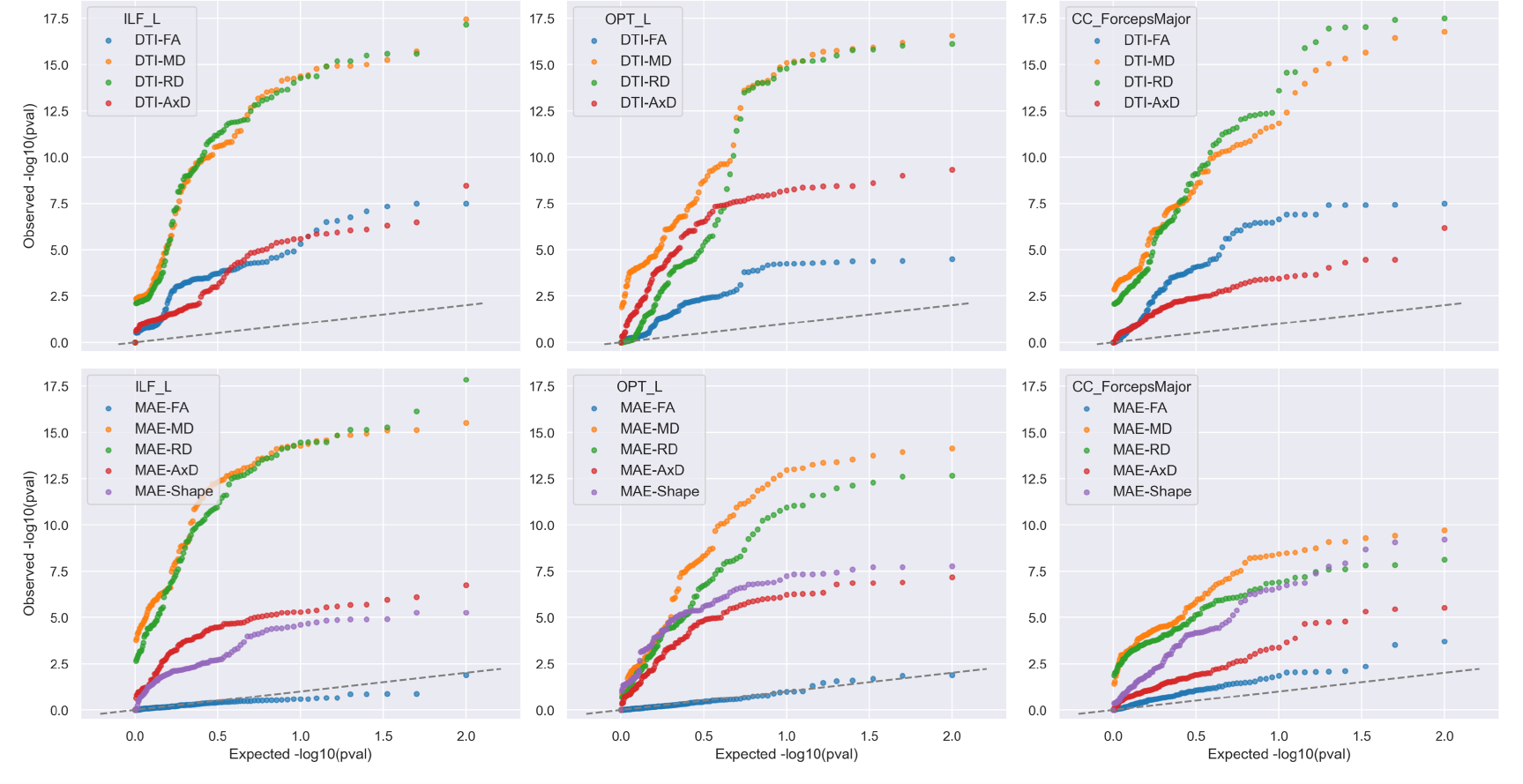
Ranking Diffusion Metrics for their Sensitivity to Dementia. Here we show a QQ plot of – log_10_(*p*) after FDR correction for dementia vs. CN group comparison in the ADNI cohort, for 4 DTI- and 5 MINT-derived diffusion measures in ILF_L, OPT_L and CC_ForcepsMajor. Microstructural metrics that are associated most strongly with dementia are typically MD and radial diffusivity (RD), shown in orange and green colors. The dashed line in these plots shows the pattern of *P* values that would be expected for null measures, i.e. brain metric that shows no association with Alzheimers disease. For some tracts, the MAE-FA is not much better than a null measure (i.e., shows no detectable association with dementia).

We selected ILF_L, OPT_L and CC_ForcepsMajor —three WM bundles from each association, projection, and commissural pathways —which showed significant dementia effects in most along-tract diffusion measures (see Section3.1). For both DTI- and MINT-derived measures, MD and RD (shown in orange and green colors) associate most strongly with dementia compared to AxD and FA for all three bundles. This is consistent with prior work examining dMRI metrics in regions of interest, where MD typically shows the strongest association with dementia. DTI-AxD can better detect dementia effects than DTI-FA in OPT_L, whereas DTI-FA outperforms DTI-AxD in CC_ForcepsMajor. Notably, MAE-FA is the weakest in detecting dementia effects in all bundles, and MAE-Shape outperforms both MAE-FA and MAE-AxD in OPT_L and CC_ForcepsMajor. These findings might indicate that metrics describing diffusion along the principle direction of axons (FA and AxD), may be more prone to crossing fibers and macrostructural changes (characterized by MAE-Shape) than MD and RD. While MAE-MD and MAE-RD remain most sensitive to dementia effects compared to other MINT-derived measures, they are less sensitive than their DTI counterparts for CC_ForcepsMajor and OPT_L. In ILF_L, on the other hand, multivariate modeling of DTI and whole-brain fiber geometry shows comparable performance in detecting dementia effects on MD and RD with univariate modeling, and can boost sensitivity for AxD measures.

## 4 DISCUSSION

### 4.1 Microstructural Brain Abnormalities in Dementia and MCI

In this study, we proposed the MINT framework for mapping micro- and macro-structural WM abnormalities using a joint multivariate model. Our novel method contrasts with the univariate approach used in existing tractometry methods such as BUAN, where metrics at each point on the tract are considered without reference to other related metrics in the rest of the brain. We investigated group differences in WM dMRI-derived metrics between dementia and MCI groups, relative to matched healthy controls, for both North American and Indian cohorts. We computed bundle profiles of the four widely-used DTI measures and five MINT-derived measures quantifying deviations from the norm, corresponding to each DTI metric and an additional shape measure. Overall, diffusivity increased in MCI and still further in dementia, with stronger abnormalities in the temporal lobe tracts and in the midline corpus callosum. Even so, we detected widespread and largely similar patterns of abnormalities across the North American and Indian cohorts with stronger statistical significance for disease effects in ADNI than NIMHANS, which is a slightly younger cohort. Of all DTI-derived metrics, MD was the most sensitive to dementia, followed by RD, AxD, and FA. Patterns of dementia and MCI effects on MD were also consistent across univariate and multivariate approaches, as well as across the two independent cohorts. This aligns with our understanding of DTI-derived metrics, in that MD is less susceptible to crossing fibers.

### 4.2 Paradoxical Patterns of Fractional Anisotropy

Notably, patterns of dementia and MCI effects on RD and AxD were similar between the corresponding DTI- and MINT-derived features except in the projection pathways, for which they exhibit a ‘fan-shaped’ pattern where the *corona radiata* and the *corpus callosum* intersect. This pattern is prominent for MAE-Shape and DTI-FA, where the proximal parts have higher DTI-FA in dementia compared to controls, contrary to what is commonly expected in WM degeneration. Areas showing higher DTI-FA in dementia and MCI largely correspond to areas with lower macrostructural anomalies characterized by MAE-Shape, with the exception of the bilateral OPT fibers. This paradoxical pattern of FA has previously been reported in dementia and MCI by ^55,56^ in regional DTI analyses with Tract-Based Spatial Statistics (TBSS) ^57^. In healthy aging, FA has also been found to have no detectable association —or even a paradoxical, positive association with age —in the projection fibers ^58–60^. Using fixel-based analysis, Han *et al*. ^61^ reported fiber-specific age associations in healthy adults and found evidence of selective WM degeneration. This ‘fan-shaped’ pattern in the projection pathways in our findings is likely due to the fiber crossings near the *corona radiata* and the *centrum semiovale*. The complex fiber configurations in the commissural and U-fibers, compared to the more homogeneous fiber organization in the deep proximal regions, may contribute to this spatially-varying pattern in the long-range projection pathways. While such fiber crossings would presumably exist in healthy controls, we found that this pattern is enhanced in dementia and MCI groups in both cohorts after accounting for age and sex effects. In a voxel-wise DTI analysis, ^62^ reported higher FA in the cingulum and fornix in preclinical AD subjects who are amyloid positive. FA is a composite measure and is not a reliable metric for interpreting fiber integrity ^11^. For similar reasons, RD and AxD are also susceptible to fiber crossings, where changes in one can lead to spurious changes in the other measure ^63^. When studying neurodegenerative diseases such as AD, where the underlying pathology may affect the biophysical properties of WM, the effects of fiber crossing can overshadow (and be confounded with) disease-related microstructural changes in FA, AxD, and RD, when considered independently. Our novel MINT framework accounts for whole-brain macrostructure using tractography in a multivariate model. This may help to disentangle intrinsic changes in DTI measures from signals affected by crossing fibers, to better understand the underlying pathology. While FA is the metric that is most correlated with alterations in WM geometry (macrostructure) —and few regions pass the statistical significance threshold compared to univariate DTI measures —MINT still identified widespread abnormalities in RD and AxD in dementia and MCI, and comparable correlation with cognitive measures compared to raw DTI values in some WM regions. Although MD is less prone to the effect of crossing fibers than other DTI metrics, we note that it is susceptible to partial volume effect. In WM regions close to the ventricles, such as the *forceps major*, CSF may contribute to changes in MD that could be attributed to atrophy instead of WM microstructural changes. We will investigate multi-compartment diffusion metrics such as free water ^64^ in the MINT framework in our future work, to better account for various sources of signal in the multivariate model.

### 4.3 Relative Strengths and Limitations of Regional DTI Metrics versus Tractometry

Although FA, RD, and AxD have major limitations in characterizing WM microstructure, they still have their place in dMRI analysis, as do simpler, region-of-interest approaches such as TBSS —as tractography often requires high *b*-values (dMRI gradient strength) and high angular resolutions and not all data can be analyzed with tractometry. Even in acquisitions with sufficient angular resolution for tractography, widely-used tractography and segmentation pipelines systematically produce false positive streamlines ^65^ For this study, we selected 30 bundles from a total of 80 bundles from the HCP-842 atlas ^8,22^. State-of-the-art fiber orientation reconstruction and tractography methods used in this study ^30,32^ can sufficiently reconstruct fibers from these bundles in single-shell dMRI acquisitions with low *b*-values ^66^. However, some key bundles, such as the fornix and the cingulum, are difficult to reconstruct using automated methods due to their unique shape and location ^67^, although the fornix is a key structure showing early changes in aging ^68^. Even among the 30 bundles we used in this study, UF and IFOF bundles are not well reconstructed in all subjects across all protocols, which may affect downstream harmonization and statistical analysis (**Supplementary Section 5**). Skeleton-based methods such as TBSS, compared to tractometry, examine a relatively small subset of the brain’s anatomy. These methods can be more computationally and memory-efficient, facilitating efficient large-scale international analyses such as those by the ENIGMA-DTI working group ^3^. FA abnormalities in neurodegenerative diseases are robustly reported in numerous studies, as are age effects across the human lifespan in normative models of vast international samples ^69,70^. While FA, RD, and AxD are not specific indices of microstructural integrity *per se*, they are still useful for group comparisons and for detecting anomalies in acquisitions where the WM tracts of interest are difficult to reconstruct using tractography.

### 4.4 Comparing Findings Across Cohorts

In our analysis of dementia and MCI effects on dMRI metrics,findings were very similar between the ADNI and NIMHANS cohorts. The difference in the apparent effect of diagnosis on brain microstructure, *β*(DX), and its statistical significance is likely due to the difference in sample sizes and the slight difference in age, as the NIMHANS sample is younger than ADNI. The pattern of dementia and MCI effects is overall more consistent for univariate DTI measures compared to MINT MAE measures. In ADNI, where the sample size is larger, MAE measures for RD and AxD are largely homogeneous along-tract compared to the smaller NIMHANS cohort. In the model fine-tuning stage, 42 CN participants from NIMHANS were analyzed compared to the 150 CN subjects from ADNI. For statistical analysis, the ADNI cohort had more CN subjects (N=226) compared to MCI (N=214) and dementia (N=69) groups, whereas the NIMHANS cohort has only 62 CN subjects compared to 89 MCI and 90 subjects with dementia. MINT may more effectively encode patterns of whole-brain fiber geometry when a larger sample of healthy controls is used in both model training and statistical analysis. As the pretrained model was fine-tuned for each cohort separately, the model may have learned to better encode fiber geometry and generalize to unseen subjects in the ADNI dataset. Although age is modeled in the linear regression model, participants from NIMHANS are younger than those in ADNI on average, and diagnostic criteria may differ across cohorts for dementia and MCI. In Figure 5, the NIMHANS cohort showed stronger dementia and MCI effects on macrostructural alterations characterized by MAE-Shape than ADNI, in projection pathway regions close to the cortex. These anomalies are also concentrated in the temporal regions of the left hemisphere association pathways in NIMHANS compared to ADNI, where more widespread alterations are detected bilaterally in the posterior and temporal association pathways. While this may be an effect of sample size, or even overfitting, we cannot discount the possibility that the impact of dementia and MCI on WM may differ across Indian and North American populations. Future work will assess how findings depend on sample sizes used for model training and testing; we will also investigate how dMRI measures relate to specific types of neuropsychological decline (e.g., language, memory, executive function), as well as amyloid and tau measures in larger populations.

### 4.5 Normative Modeling and Harmonization for Tractometry

Generative models such as the VAE are well-suited for normative modeling and anomaly detection, as they can model the joint data distribution of multiple complementary measures (even high dimensional data such as bundle profiles and even 3D MR images). They can more fully model within-group variation in brain structures compared to the traditional case-control schema. Normative models based on VAEs can be used for single subject anomaly detection ^71^ as well as group difference testing, as in this study. Compared to traditional normative models such as those based on the Generalized Additive Model for Location, Scale, and Shape (GAMLSS) ^72^ and Hierarchical Bayesian Regression (HBR) ^73^, autoencoder-based normative models have different model formulations, harmonization, training, and inference procedures. Normative models in neuroimaging often estimate the normative range for a regional brain measure given covariates from a sample of healthy controls. However, they are typically univariate models - where the measure being predicted is a single measure from one brain region - and cannot model spatial correlations between regions ^74^ or covariance between multiple measures. By contrast, VAEs can learn complex joint distributions of high-dimensional data, as in this study, where we used them to jointly model whole-brain WM macro- and micro-structure. Recent extensions of VAEs to latent diffusion models (LDMs) show the power of the generative modeling approach, as they can generate entire realistic brain images with prescribed age, sex, and even disease characteristics ^75,76^. The VAE architecture may be designed to more effectively model a large feature space: the sequential dependency along each streamline is modeled with 1D convolution, and the streamlines’ flip-invariance can be enforced using random flipping for data augmentation.

In terms of data harmonization, the hierarchical formulation in HBR can model the mean and variance structure for each site directly without using ComBat type approaches and produce a site-agnostic z-score. Among autoencoder approaches, a conditional VAE can be used to learn representations in the latent space —that are invariant to site —and produce images that appear as though they were acquired at a different site, or with a different scanning protocol ^77^. To use CVAE in a normative model setting, ^78^ proposed a deviation metric defined in the latent space without the effect of confounds such as age or intracranial volume (ICV). However, there is a trade-off between deviation metrics with high specificity and direct confound removal in CVAE models. Deviation metrics defined for latent variables often produce one measure per sample (3D MRI volume or streamline in MINT). In contrast, metrics defined from reconstruction error can produce one measure for every unit in the sample (a voxel in a 3D volume or point on a streamline). As the latent representations and decoder reconstruction are ‘confound-free’ and the original input is not, reconstruction-based metrics should not be used to remove confounds or site effects in CVAE-based normative models.

In MINT, we opted for a VAE model with ComBat harmonization ^17^ instead of a CVAE to obtain fine-scale mapping of along-tract WM deviations. We found that ComBat can be successfully applied to a deviation metric such as MAE, and the fixed effect coefficient for diagnosis was similar before and after ComBat (see **Supplementary Section 5**). Although MAE is non-negative and right skewed, ComBat can be applied to non-Gaussian distributions ^79^. However, ComBat may introduce artifacts and inflate group differences ^80^ in samples where the site is confounded with variables of interest (e.g., if inclusion or exclusion criteria differ at any sites). In the future, we will experiment with other variations of ComBat, such as ComBat-GAM ^47^ and CovBat ^81^, to model non-linear effects of covariates. In related recent work, the eHarmonize software ^70^ was created to harmonize new datasets to lifespan reference curves created from regional DTI measures in large multi-site samples. It supports training, testing, and quality control using a ComBat-type method, and may also be adapted to tractometry analysis.

### 4.6 Modeling Tractography Data Efficiently

Efficiency is essential when working with densely sampled tractography data, which may ultimately include data from thousands of subjects. We designed a lightweight VAE model for MINT with less than half a million parameters; many design choices are described in Section 2.3.2. The training data used in this study (including those for pre-training and fine-tuning) exceed 500GB in size, but the memory requirement for MINT is low as streamlines are loaded in batches for training and inference.

MINT uses transfer learning for more efficient adaptation to new datasets. Transfer learning can be implemented using supervised, unsupervised, or self-supervised pre-training approaches and can increase downstream task performance ^82^. Pretraining can leverage other datasets, and significantly speed up convergence when applied to a new dataset. ^83^ compared various pre-training strategies and their sample size requirement in an AD classification task from T1-weighted structural MRI. Contrastive learning performed best, even when the pre-training sample size was small. Contrastive learning has been applied to tractography data for bundle segmentation ^40,84^ and predictive modeling ^85^, and could also be investigated in the MINT framework.

Aside from pre-training strategies, the choice of pre-training data, sample size, and features are also crucial for model generalization. We discussed in Section 4.3 that not all WM tracts of interest can be reconstructed with tractography in clinical dMRI acquisitions. We investigated the micro- and macro-structural properties of deep WM tracts in this study, but one limitation is that we did not examine U-fibers - the small tracts that connect adjacent cortical gyri. These would be valuable targets for future work, not least because the accumulation of cortical amyloid appears to influence dMRI signals in the cortical gray matter and might, therefore, affect the short association fibers between adjacent gyri of the brain. Also, since the RecoBundles method extracts each bundle separately, the same streamline from a whole-brain tractogram can be segmented for multiple bundles, mainly if they lie in close proximity and share similar structural patterns. Depending on the choice of atlas, some bundles may be over-represented in the training data. As the MINT framework uses a streamline-based model, it may be more desirable to use whole-brain tractograms directly, especially in the pre-training stage. However, they often contain many false positives that may hinder model training ^10^, and it is important to evaluate the trade-off between maximizing whole-brain information to better model fiber crossings and reducing false positives. In this study, we selected the TractoInferno dataset as it is multisite and uses single-shell acquisitions. It is also publicly available, preprocessed, and carefully quality-controlled. However, in the future, we will assess tractography data from multi-shell acquisitions such as the Human Connectome Project (HCP) ^86^ in the pre-training stage to better learn the organization of complex fiber populations, and apply tractogram filtering techniques such as SIFT/SIFT2^87,88^, COMMIT/COMMIT2^89,90^ and FINTA ^38^ to reduce the proportion of false positive streamlines. Additionally, diffusion MRI measures derived from multi-shell data, such as neurite orientation dispersion and density imaging (NODDI) ^64^, diffusion kurtosis imaging (DKI) ^91^, and MAP-MRI ^92^ may provide a richer feature set and may detect AD-related abnormalities with greater sensitivity and specificity ^4,93,94^.

## ACKNOWLEDGMENTS

Algorithm development for this project was supported by the NIA under grant R01 AG057892. The NIMHANS data collection was support by NIH grant R01 AG060610 and the Department of Science and Technology, Govt. of India, grant nos. DST-SR/CSI/73/2011 (G); DST-SR/CSI/70/2011 (G); and DST/CSRI/2017/249 (G). Data collection and sharing for the Alzheimer’s Disease Neuroimaging Initiative (ADNI) is funded by the National Institute on Aging (National Institutes of Health Grant U19 AG024904). The grantee organization is the Northern California Institute for Research and Education. In the past, ADNI has also received funding from the National Institute of Biomedical Imaging and Bioengineering, the Canadian Institutes of Health Research, and private sector contributions through the Foundation for the National Institutes of Health (FNIH) including generous contributions from the following: AbbVie, Alzheimers Association; Alzheimers Drug Discovery Foundation; Araclon Biotech; BioClinica, Inc.; Biogen; Bristol-Myers Squibb Company; CereSpir, Inc.; Cogstate; Eisai Inc.; Elan Pharmaceuticals, Inc.; Eli Lilly and Company; EuroImmun; F. Hoffmann-La Roche Ltd and its affiliated company Genentech, Inc.; Fujirebio; GE Healthcare; IXICO Ltd.; Janssen Alzheimer Immunotherapy Research & Development, LLC.; Johnson & Johnson Pharmaceutical Research & Development LLC.; Lumosity; Lundbeck; Merck & Co., Inc.; Meso Scale Diagnostics, LLC.; NeuroRx Research; Neurotrack Technologies; Novartis Pharmaceuticals Corporation; Pfizer Inc.; Piramal Imaging; Servier; Takeda Pharmaceutical Company; and Transition Therapeutics.

We thank the research study participants and their families whose contributions made this work possible. We also thank the DIPY, FURY, and pytorch-lightning contributors and community, whose tools facilitated the analyses in this study.

## Footnotes

‡ For a summary of related work on methods for dMRI analysis, please see **Supplementary Material Section 1**.

§ S, P, and GE denote Siemens, Philips, and General Electric MRI scanners, and the number in the abbreviation denotes the number of diffusion-weighted gradients in each protocol; if other factors are constant, higher numbers of gradients tend to give better signal-to-noise ratio in dMRI.

¶ Along-tract *β*(DX) for DTI-RD, MAE-RD, DTI-AxD and MAE-AxD are available in the **Supplementary Figures S1-2**. The corresponding along-tract – log_10_ *P* values after FDR correction for all 9 measures are available in **Supplementary Figures S3-7**

