## Supplementary Materials for "Microstructural Mapping of Neural Pathways in Alzheimer’s Disease using Macrostructure-Informed Normative Tractometry"

### 1 Related Methods for Diffusion MRI Analysis

Widely used methods to quantify WM microstructural differences between groups include voxel-based, fixel-based and tractometry approaches. *Voxel-based methods* first register voxel-based scalar maps — such as those derived from DTI —to an atlas space, and then conduct univariate tests at the voxel or region-of-interest (ROI) level. Tract-based spatial statistics (TBSS) [1] was proposed to better align WM structures across subjects, by extracting the WM skeleton from fractional anisotropy (FA) maps, and summarizing white matter metrics in regions of the skeleton. When studying diffusion MRI, we also have to consider the special nature of WM structure: modeling how signals propagate along the long-range WM fibers is more faithful to the underlying anatomy than modeling signals in an isotropic neighborhood. Fixel-based analyses, such as connectivity-based fixel enhancement (CFE) [2], quantify microstructural properties in specific fiber populations within a voxel to better resolve crossing fibers, with special procedures for spatial normalization and statistics. Another family of methods - *tractometry* [3, 4, 5] - creates *bundle profiles* or *along-tract profiles* [6, 7, 8] by projecting local measures of tissue microstructure, onto 3D white matter bundles reconstructed from tractography. Automatic Fiber Quantification (AFQ) [6] computes a mean bundle profile of microstructural measures —where the measure from each streamline (3D curve) in a bundle is weighted by distance to the bundle core. Bundle Analytics (BUAN) [7] uses the full structure of a bundle, yielding smoother along-tract metrics. BUAN has identified microstructural abnormalities in Parkinson's disease (PD), Alzheimer's disease (AD) [9] and bipolar disorder [10, 11] compared to matched controls. A more recent tractometry approach, Medial Tract Analysis (MeTA) [12] computes the core volume of a bundle around the medial surface for microstructural mapping to improve the reliability of bundle profiles. Many tractometry methods analyze segments/nodes along the length of bundles independently without taking into account neighboring information of points on a streamline. [13] proposed to model streamlines as functions to take into account neighboring information of points on the streamline.

Tractography, and by extension tractometry, demand substantial memory and computational resources, making it challenging to scale analyses to large cohorts, but the large volume of data can potentially be analyzed using deep learning methods. The unique format of tractography data is important to consider when adapting deep learning model architectures used in other domains. Point-cloud based networks have been applied to tractography data for bundle segmentation [14] and predictive modeling [15, 16]. These models represent bundles as point clouds —as sets of 3D points, instead of a collection of streamlines —which are sets of ordered 3D point sequences. Without using a voxel grid, deep learning methods, such as PointNet [17], can encode point cloud data represented with 3D coordinates using operations that are invariant to permutations or arbitrary ordering of the data, such as fully connected layers and global pooling. These methods can extract bundle-level information, but they do not capture the dependency of neighboring points on a single streamline —an important source of macrostructural information derived from fiber tracking. Additionally, given the large number of streamlines per bundle, data reductions may be required to use a larger batch size during model training. In our prior work [18], we showed that a variational autoencoder (VAE) with 1D convolutional layers [19] can embed streamlines into a compact latent space and be used to detect structural anomalies in AD, and generate synthetic bundles via generative sampling.

Autoencoder-based architectures can also be used in normative models [20, 21, 22], to encode statistical distributions of features from a reference population. Deviations from the norm can be quantified and used in downstream analysis for group difference testing or mapping individual anomalies [23]. After training an autoencoder, data from the patient group is passed through the network at inference time, and the reconstruction error can be used for anomaly detection, or for group statistical comparisons. [24] first proposed *normative tractometry* to localize along-tract microstructural anomalies using autoencoders, revealing abnormalities in single subjects with genetic copy number variants (CNVs), epilepsy and schizophrenia. In our previous study using tractography data from the Alzheimer's Disease Neuroimaging Initiative (ADNI), our ConvVAE-based model identified 6 WM tracts with along-tract macrostructural anomalies in AD [25]. Using traditional machine learning methods, normative models have been used to quantify deviations from the normal range of variation in brain morphometry [26, 27] and functional network metrics [23] in large multi-site samples. Recent efforts have also studied age effects on brain microstructure over the human lifespan, producing normative charts for the primary microstructural metrics based on data from over 40,000 healthy individuals [28, 29, 30]. When merging multisite data to increase sample sizes, or to test the generalizability of the effects to different populations, the variability introduced by site (or scanning protocol) can strongly impact statistical analyses. Sources of multi-site variability include scanners from different vendors (e.g., GE, Siemens, or Philips), different acquisition protocols, and inclusion criteria which can affect sample characteristics [31]. Diffusion MRI is especially susceptible to protocol effects, although they can be carefully modeled using harmonization techniques [32, 33, 34]. A widely used method for data harmonization in neuroimaging, ComBat [35, 36] was recently adapted and extended to along-tract metrics, for use with the BUAN tractometry pipeline [37].

### 2 Atlas Bundles

Table 1: Thirty bundles from HCP842 atlas [38] used in bundle segmentation.

| Abbreviation | Full Name | Category |
|---|---|---|
| AF_L | Left Arcuate Fasciculus | Association |
| AF_R | Right Arcuate Fasciculus | Association |
| EMC_L | Left Extreme Capsule | Association |
| EMC_R | Right Extrame Capsule | Association |
| IFOF_L | Left Inferior Fronto-occipital Fasciculus | Association |
| IFOF_R | Right Inferior Fronto-occipital Fasciculus | Association |
| ILF_L | Left Inferior Longitudinal Fasciculus | Association |
| ILF_R | Right Inferior Longitudinal Fasciculus | Association |
| MdLF_L | Left Middle Longitudinal Fasciculus | Association |
| MdLF_R | Right Middle Longitudinal Fasciculus | Association |
| UF_L | Left Uncinate Fasciculus | Association |
| UF_R | Right Uncinate Fasciculus | Association |
| CST_L | Left Corticospinal Tract | Projection |
| CST_R | Right Corticospinal Tract | Projection |
| FPT_L | Left Frontopontine Tract | Projection |
| FPT_R | Right Frontopontine Tract | Projection |
| OPT_L | Left Occipito Pontine Tract | Projection |
| OPT_R | Right Occipito Pontine Tract | Projection |
| OR_L | Left Optic Radiation | Projection |
| OR_R | Right Optic Radiation | Projection |
| CC_ForcepsMajor | Corpus Callosum Major | Commissural |
| CC_ForcepsMinor | Corpus Callosum Minor | Commissural |
| CCMid | Corpus Callosum Mid | Commissural |
| MLF_L | Left Medial Longitudinal fasciculus | Brainstem |
| MLF_R | Right Medial Longitudinal fasciculus | Brainstem |
| ML_L | Left Medial Lemniscus | Brainstem |
| ML_R | Right Medial Lemniscus | Brainstem |
| STT_L | Left Spinothalamic Tract | Brainstem |
| STT_R | Right Spinothalamic Tract | Brainstem |
| V | Vermis | Cerebellum |

### 3 Association of Diffusion Measures with Diagnostic Groups

#### 3.1 MCI and AD Effects on Radial Diffusivity

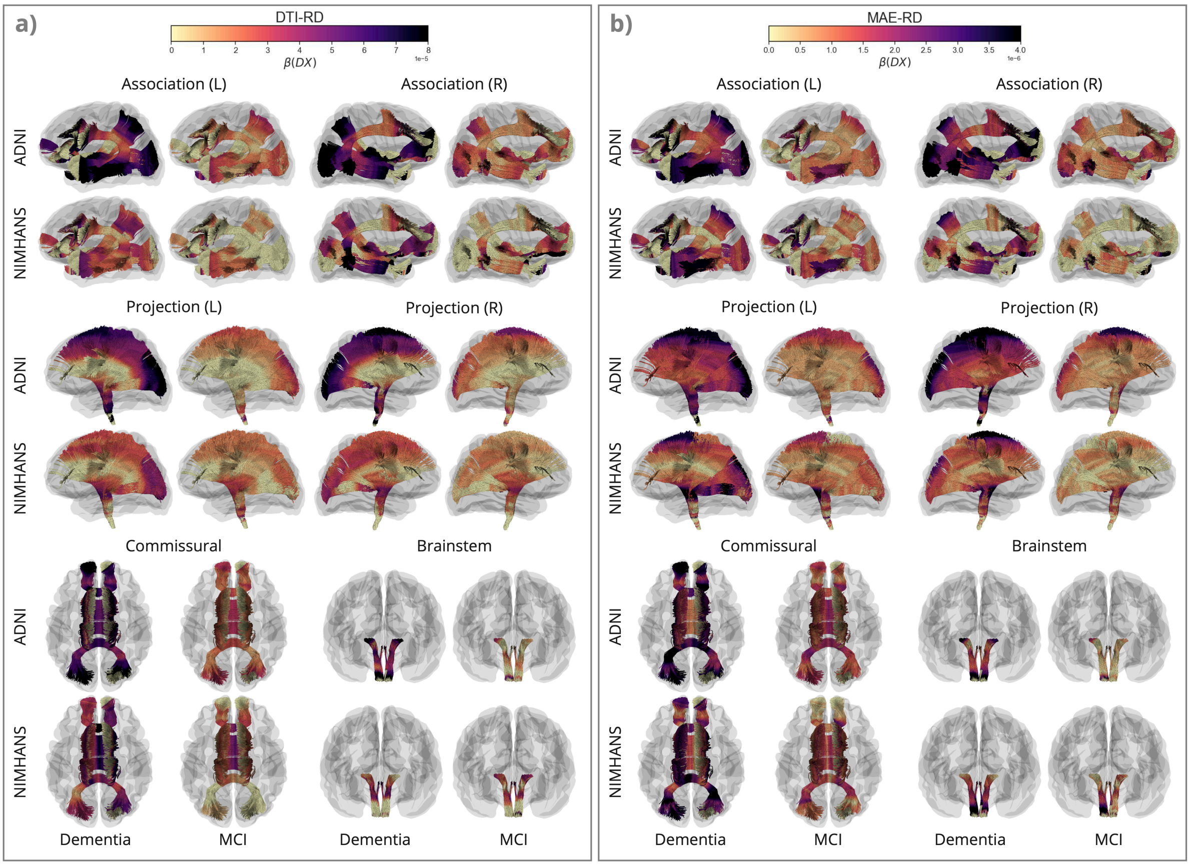

Figure 1: Along-tract $\beta$(DX) for DTI-RD and MAE-RD, for AD vs. CN and MCI vs. CN in the ADNI and NIMHANS cohorts, categorized by WM pathways.

#### 3.2 MCI and AD Effects on Axial Diffusivity

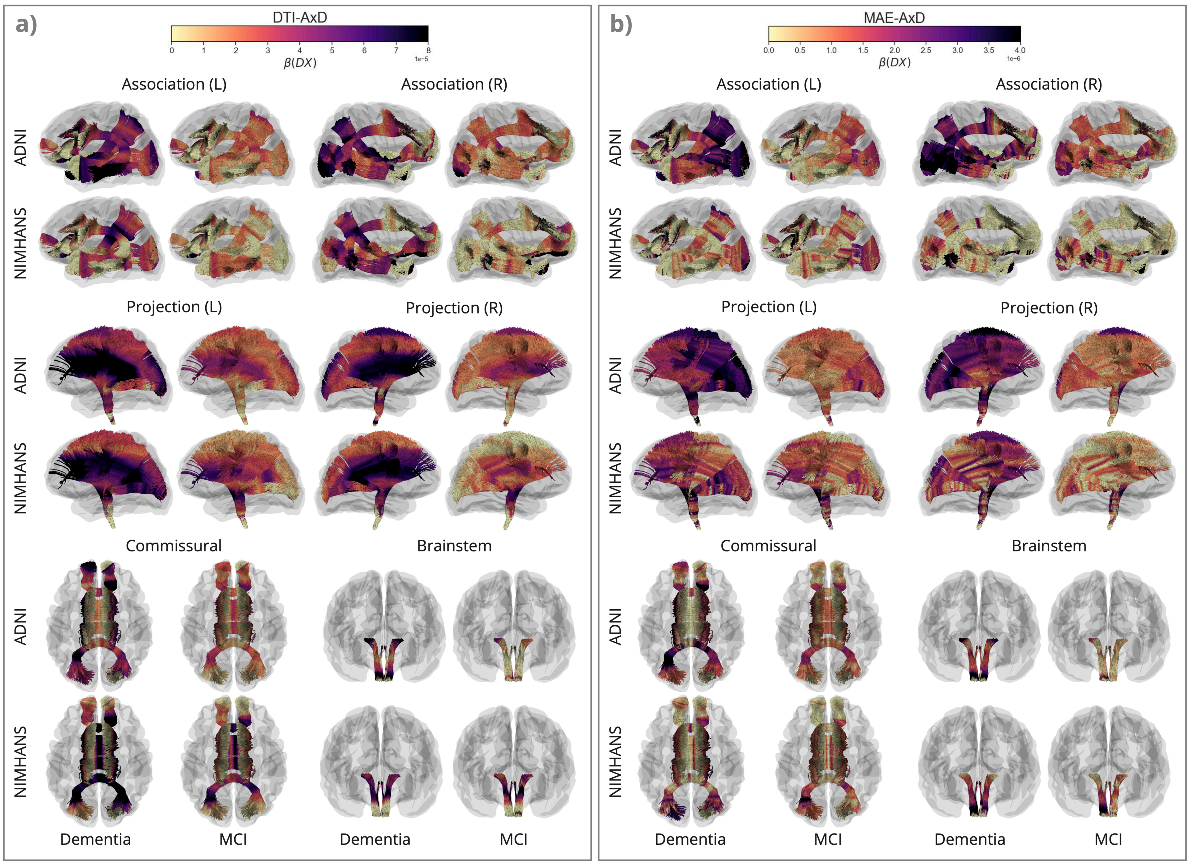

Figure 2: Along-tract $\beta$(DX) for DTI-AxD and MAE-AxD, for AD vs. CN and MCI vs. CN in the ADNI and NIMHANS cohorts, categorized by WM pathways.

### 4 Statistical Significance of Association with Diagnostic Groups

#### 4.1 Statistical Significance of MCI and AD Effects on Mean Diffusivity

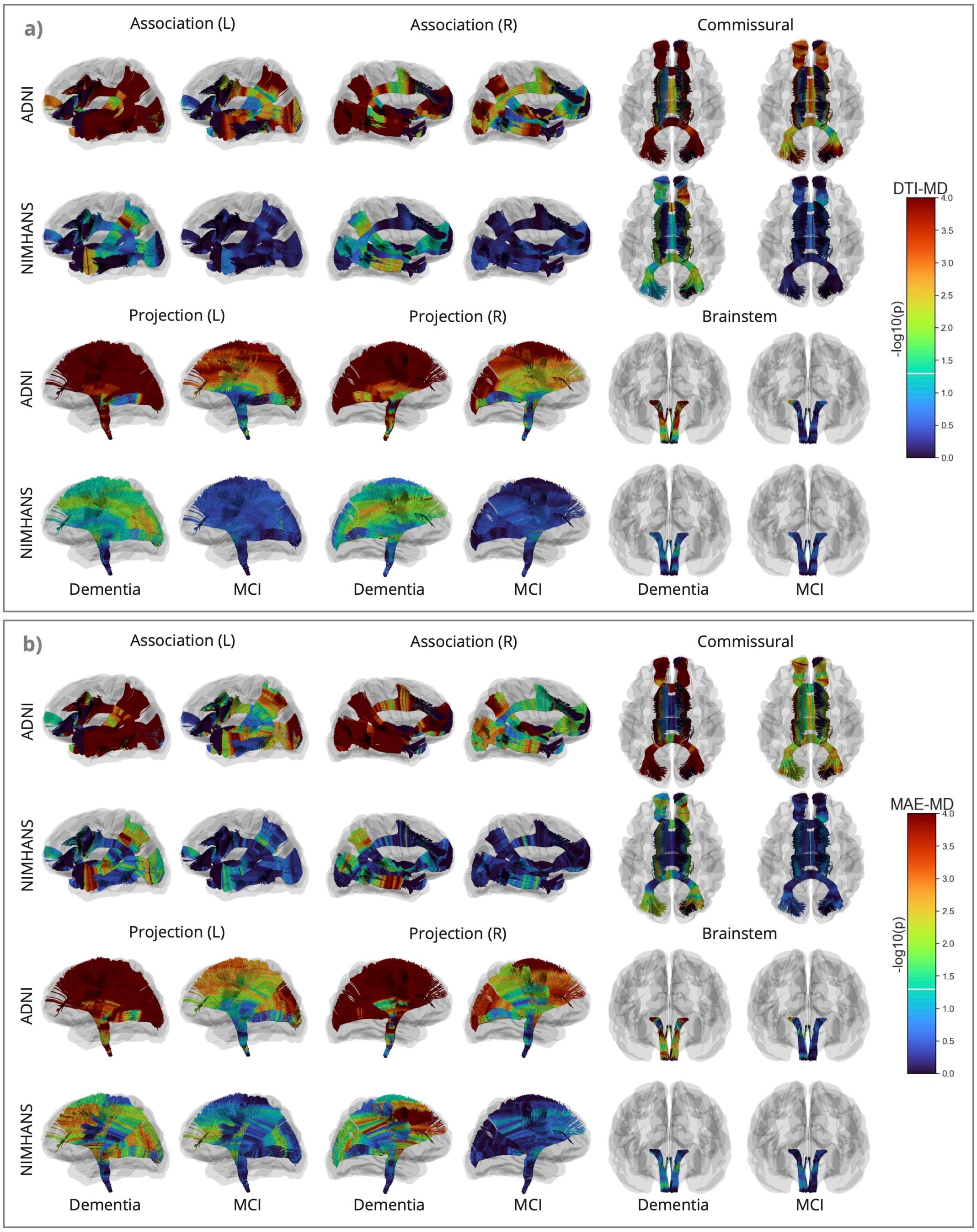

Figure 3: Along-tract $-\log_{10}(p)$ after FDR correction for MAE-MD and DTI-MD, for AD vs. CN and MCI vs. CN in the ADNI and NIMHANS cohorts, categorized by WM pathways.

#### 4.2 Statistical Significance of MCI and AD Effects on Radial Diffusivity

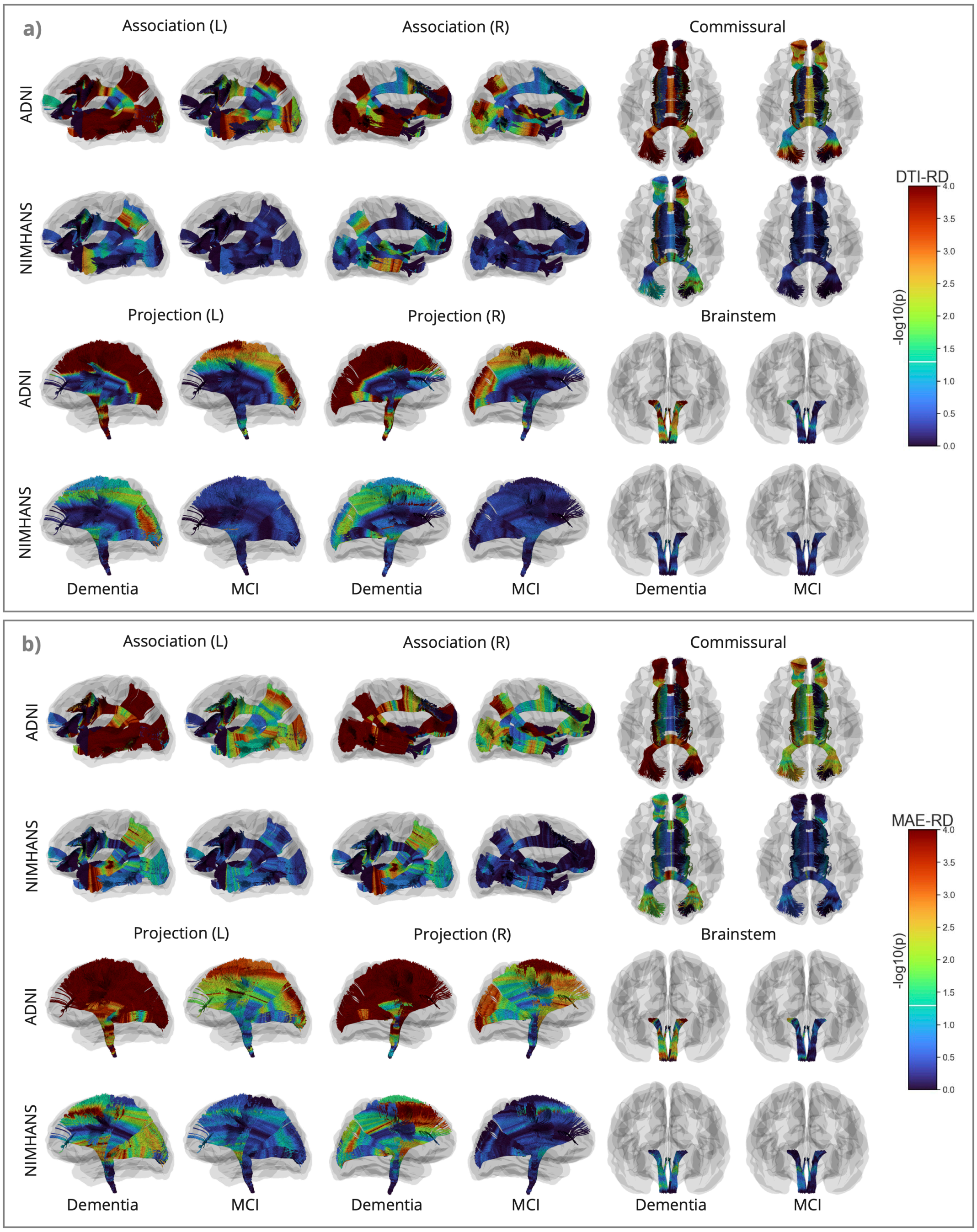

Figure 4: Along-tract $-\log_{10}(p)$ after FDR correction for MAE-RD and DTI-RD, for AD vs. CN and MCI vs. CN in the ADNI and NIMHANS cohorts, categorized by WM pathways.

##### 4.3 Statistical Significance of MCI and AD Effects on Axial Diffusivity

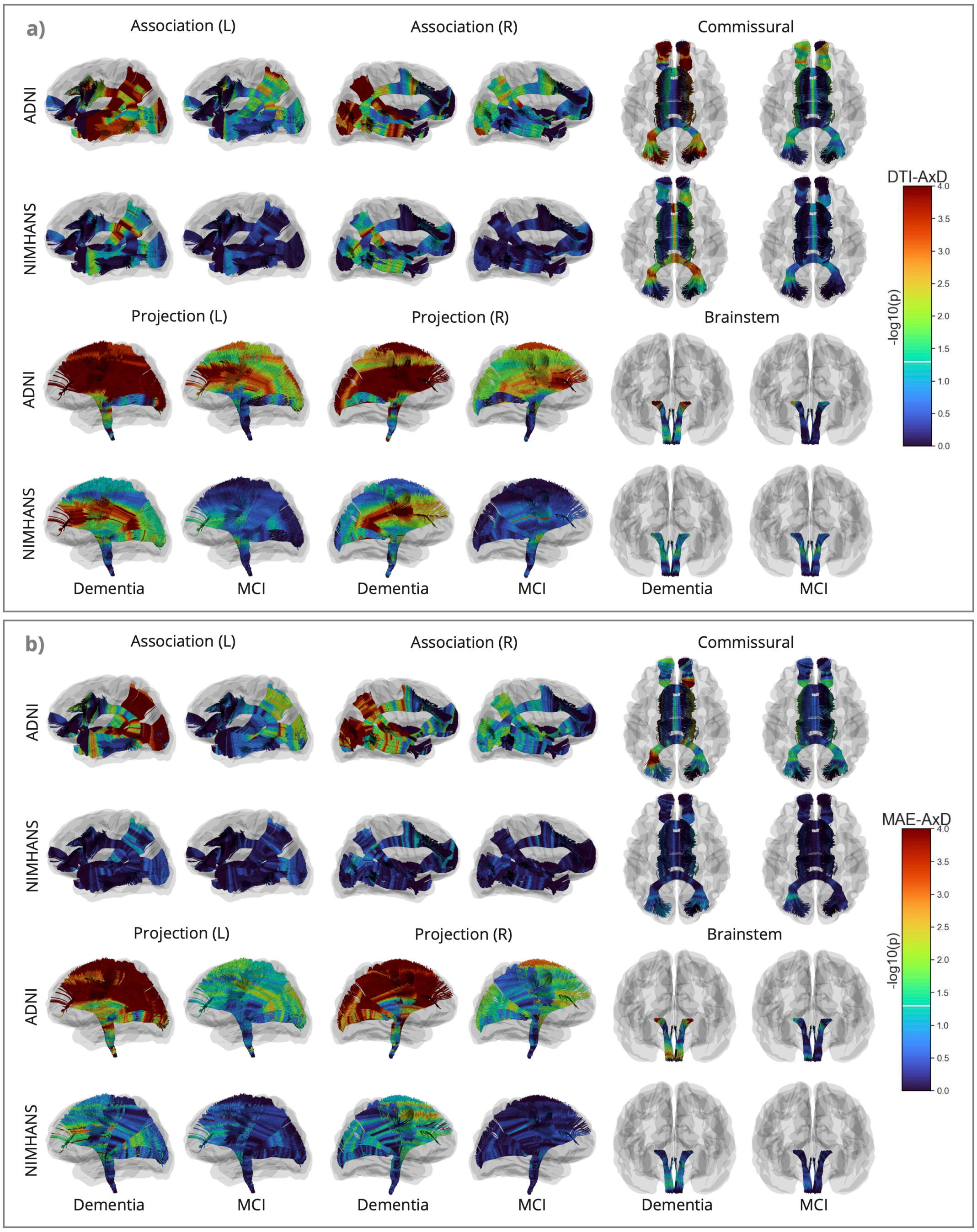

Figure 5: Along-tract $-\log_{10}(p)$ after FDR correction for MAE-AxD and DTI-AxD, for AD vs. CN and MCI vs. CN in the ADNI and NIMHANS cohorts, categorized by WM pathways.

##### 4.4 Statistical Significance of MCI and AD Effects on Fractional Anisotropy

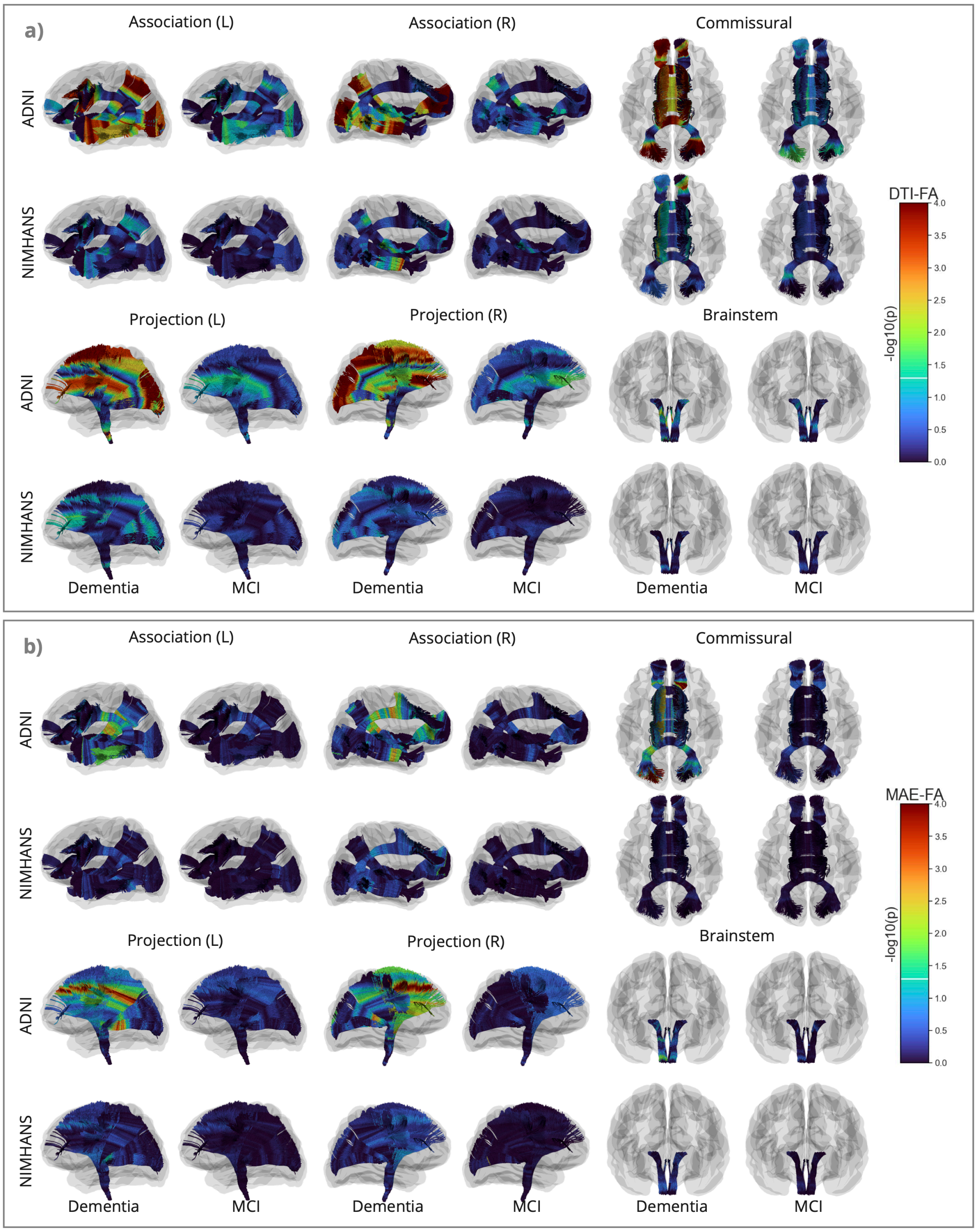

Figure 6: Along-tract $-\log_{10}(p)$ after FDR correction for MAE-FA and DTI-FA, for AD vs. CN and MCI vs. CN in the ADNI and NIMHANS cohorts, categorized by WM pathways.

#### 4.5 Statistical Significance of MCI and AD Effects on Shape Abnormalties

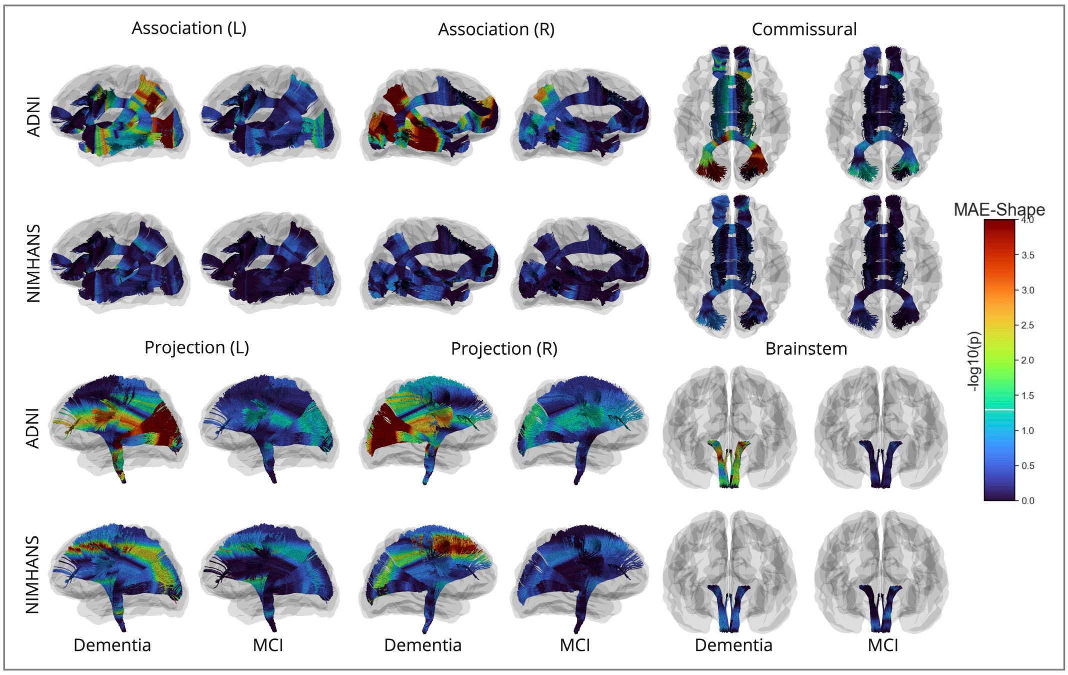

Figure 7: Along-tract $-\log_{10}(p)$ after FDR correction for MAE-Shape, for AD vs. CN and MCI vs. CN in the ADNI and NIMHANS cohorts, categorized by WM pathways.

### 5 Harmonization Evaluation of DTI- and MINT-Derived Measures

##### 5.0.1 Mean Bundle Profiles of Diffusion Indices Before and After Harmonization

All 9 DTI and MAE measures show better alignment and the trends are well preserved after harmonization for most bundles. Protocol effects vary across bundles for all measures, but their effects on MAE measures are more consistent across bundles compared to the corresponding DTI measures. The overall trends of protocol effects also differ across metrics for the same bundle, except between 3 diffusivity measures (MD, RD and AxD). These results show that harmonization applied for each bundle and measure is appropriate. Interestingly, MAE-Shape is different across cohorts but similar between protocols within the same cohort. This is likely due to model fine-tuning applied separately for each cohort, or different preprocessing pipelines. The NIMHANS Siemens protocol shows the greatest protocol difference when compared to the Philips protocol from the same cohort or the ADNI protocols. Out of all bundles, the uncinate fasciculus (UF) and inferior fronto-occipital fasciculus (IFOF) bundles show the least alignment after ComBat harmonization. This may be due to inconsistent quality of tractography and segmentation of these bundles across all protocols.

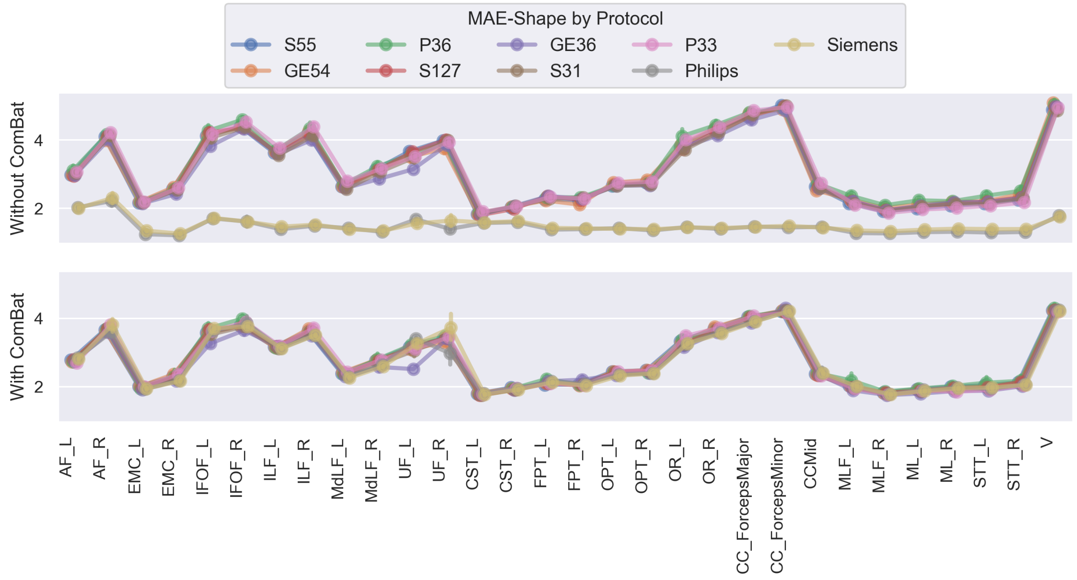

Figure 8: MAE-Shape averaged per bundle, before and after ComBat harmonization, grouped by scanning protocols from both ADNI and NIMHAN.

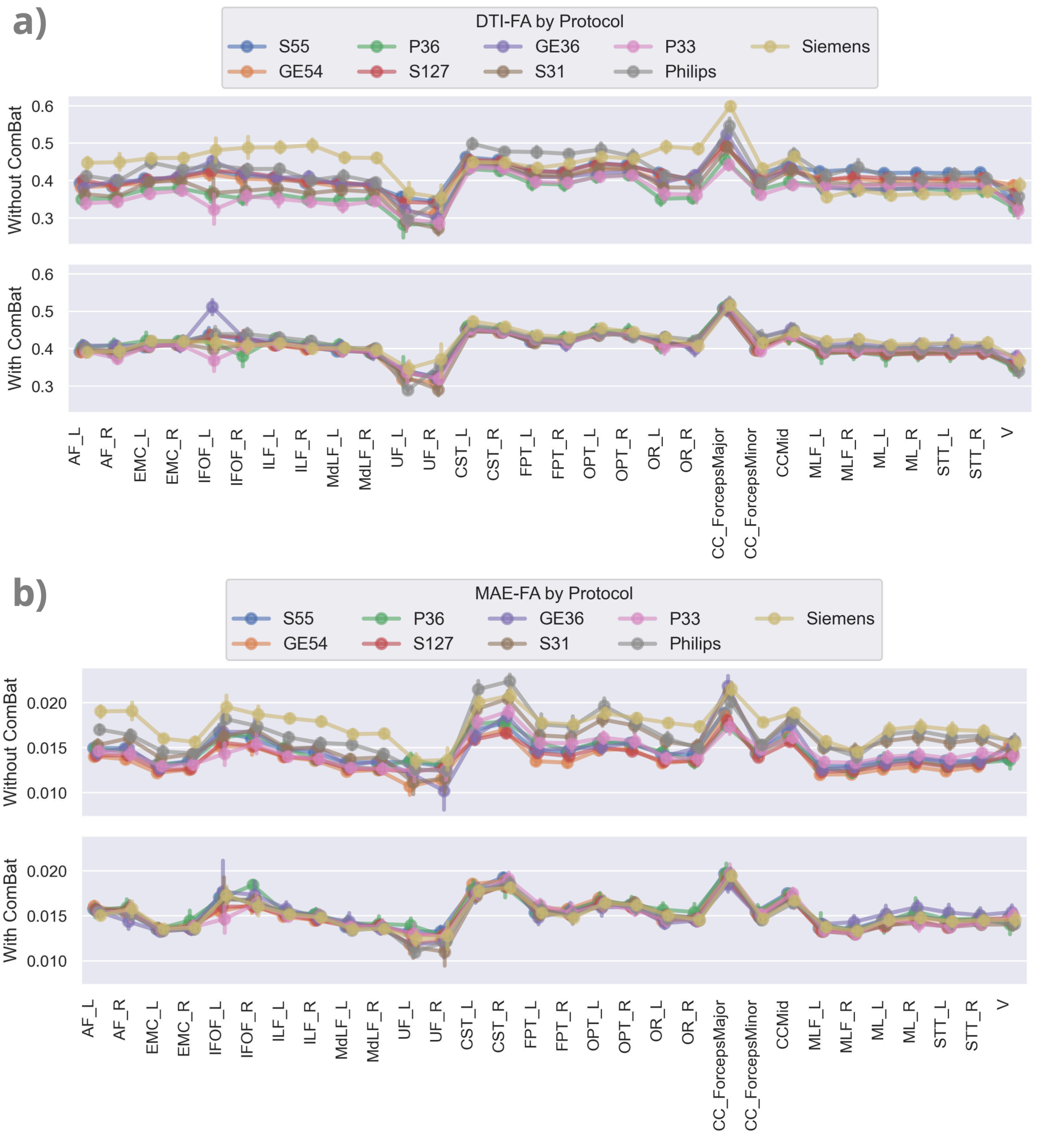

Figure 9: DTI-FA (a) and MAE-FA (b) averaged per bundle, before and after ComBat harmonization, grouped by scanning protocols from both ADNI and NIMHAN.

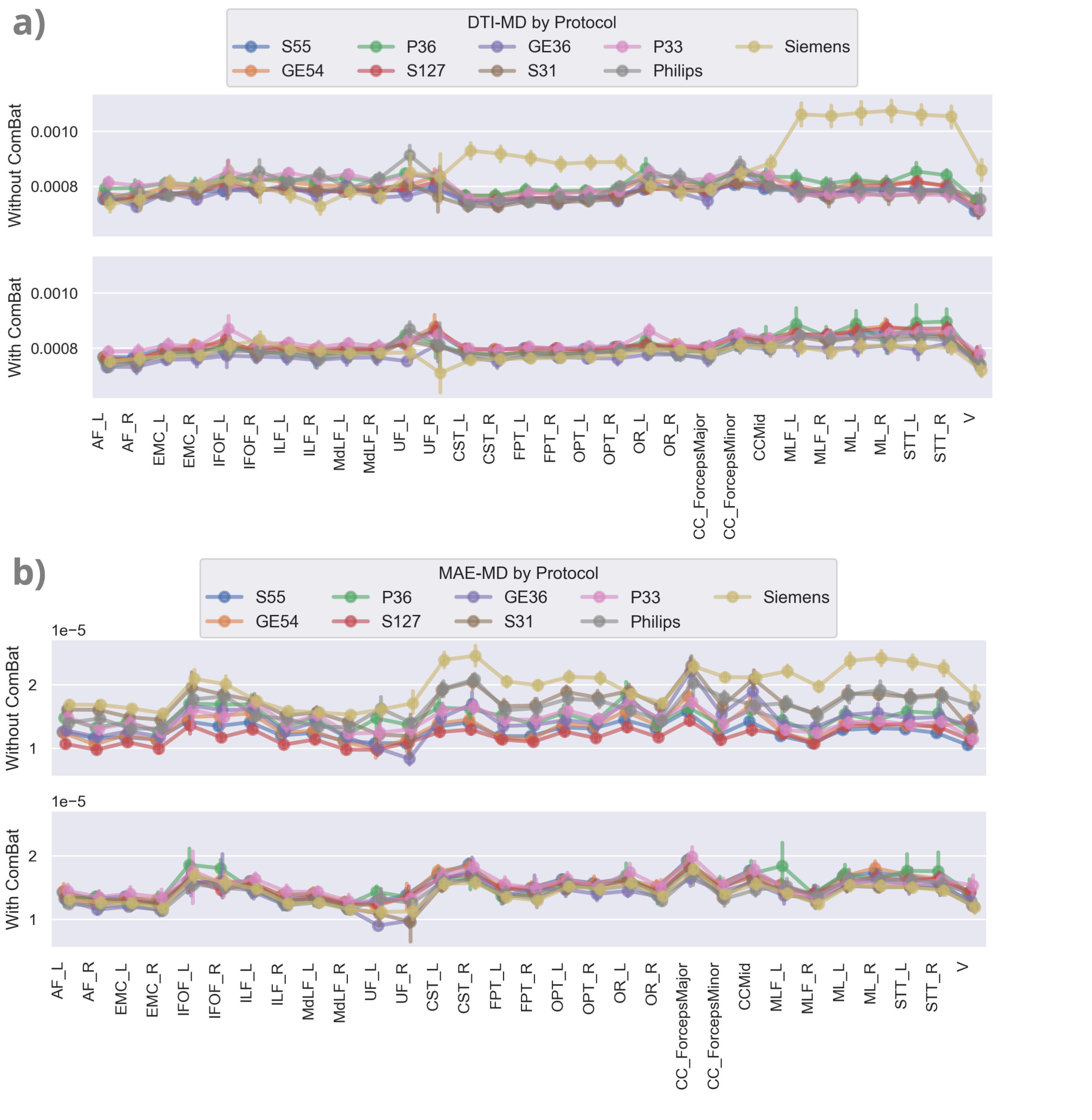

Figure 10: DTI-MD (a) and MAE-MD (b) averaged per bundle, before and after ComBat harmonization, grouped by scanning protocols from both ADNI and NIMHAN.

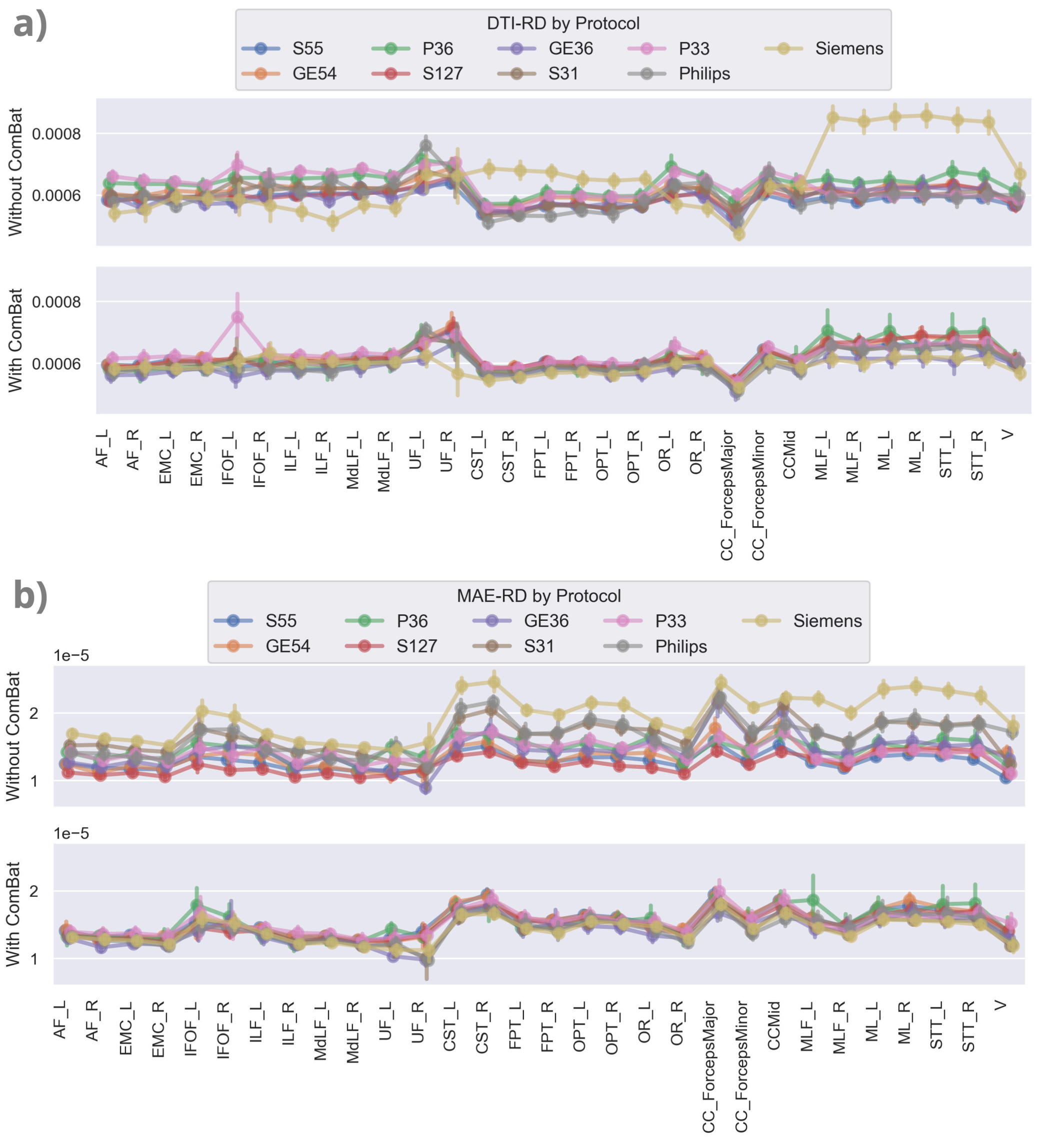

Figure 11: DTI-RD (a) and MAE-RD (b) averaged per bundle, before and after ComBat harmonization, grouped by scanning protocols from both ADNI and NIMHAN.

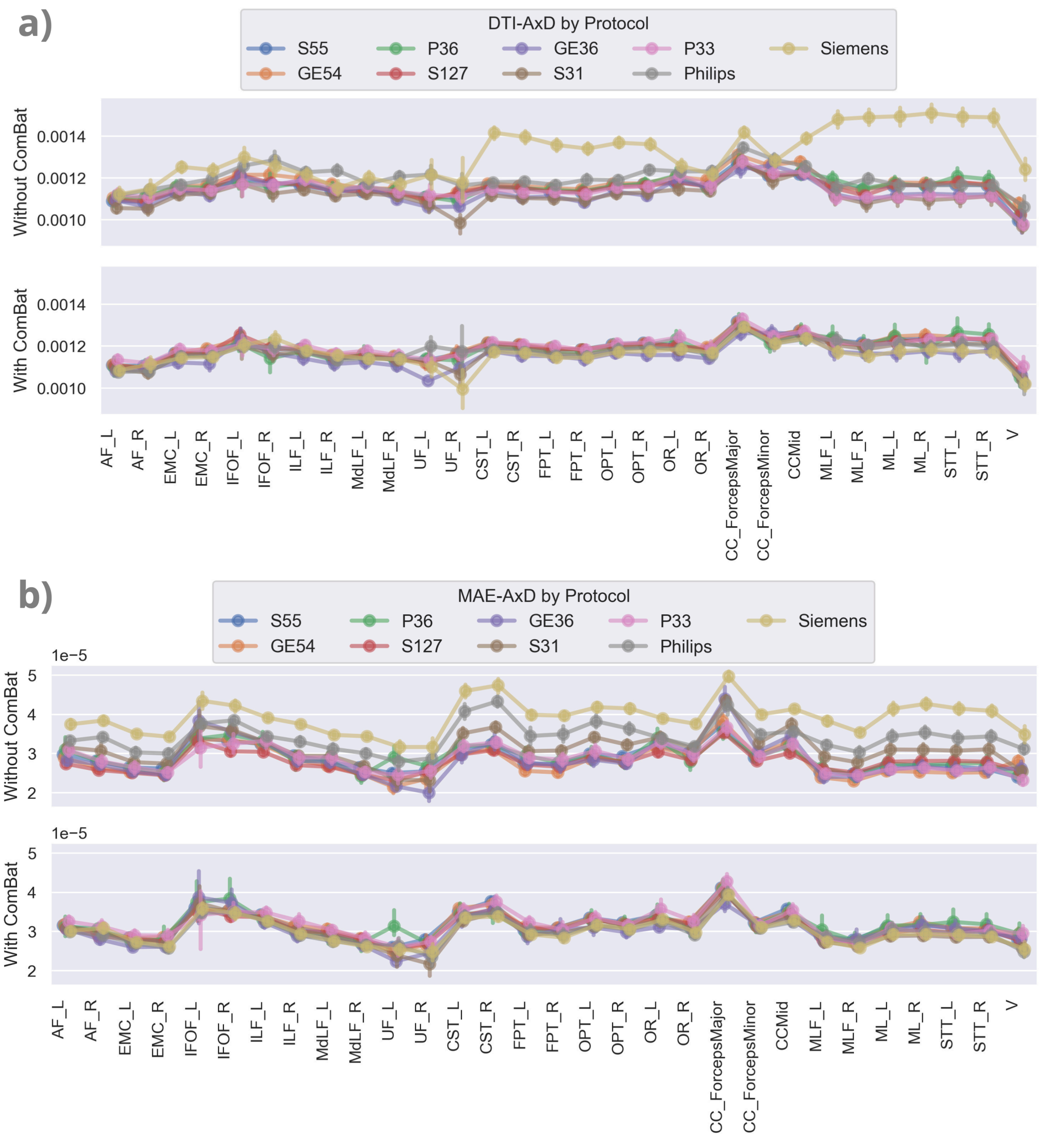

Figure 12: DTI-AxD (a) and MAE-AxD (b) averaged per bundle, before and after ComBat harmonization, grouped by scanning protocols from both ADNI and NIMHAN.

##### 5.0.2 Fixed Effect Coefficients Before and After Harmonization

To evaluate how ComBat influences statistical analysis, we compared the fixed effect coefficient for diagnosis $\beta$(DX) from the linear regression models before and after using ComBat to harmonize the data. If ComBat was successful in removing site effects while preserving biological variability, $\beta$ should remain the same before and after harmonization [39]. In Figure 13 and 14, we plot the $\beta$(DX) from the AD vs. CN comparison per bundle, for MAE features for both ADNI and NIMHANS, with and without ComBat. Overall, $\beta$(DX) are largely the same for most bundles across all MAE metrics in both cohorts, and the trends are preserved before and after ComBat. $\beta$(DX) is more consistent in ADNI than NIMHANS, except we see a large AD effect on all MAE microstructural metrics in FPT_R, AF_R, and a decreased effect on MAE-Shape in the brainstem bundles after ComBat. In the NIMHANS cohort, we see larger differences of $\beta$(DX) before or after harmonization compared to the ADNI cohort, primarily in the brainstem bundles, where the AD effect on all MAE metrics except MAE-FA is larger after ComBat. Considering that ComBat did not align the UF mean bundle profiles very well, as noted in Section 5.0.1, we also see that $\beta$(DX) in these bundles exhibits larger MAE —for all metrics —after ComBat.

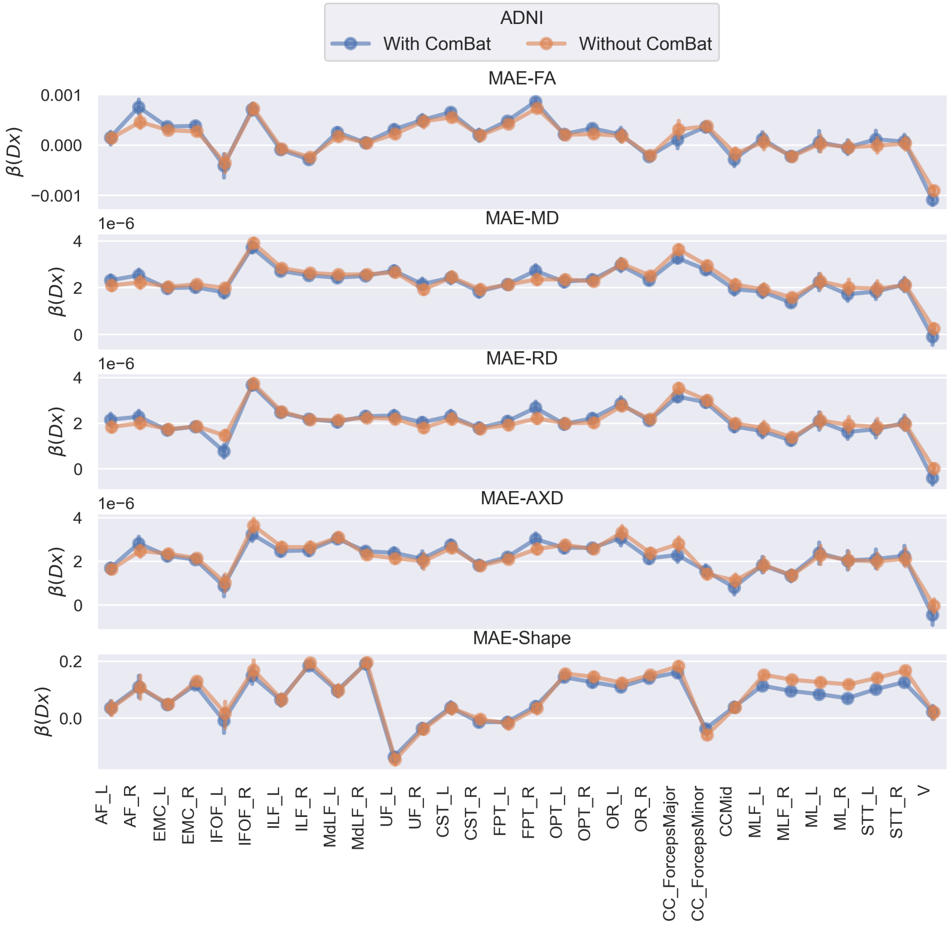

Figure 13: Fixed effect coefficient of diagnosis $\beta$(**DX**) before and after ComBat harmonization for MAE measures calculated from subjects in the ADNI cohort.

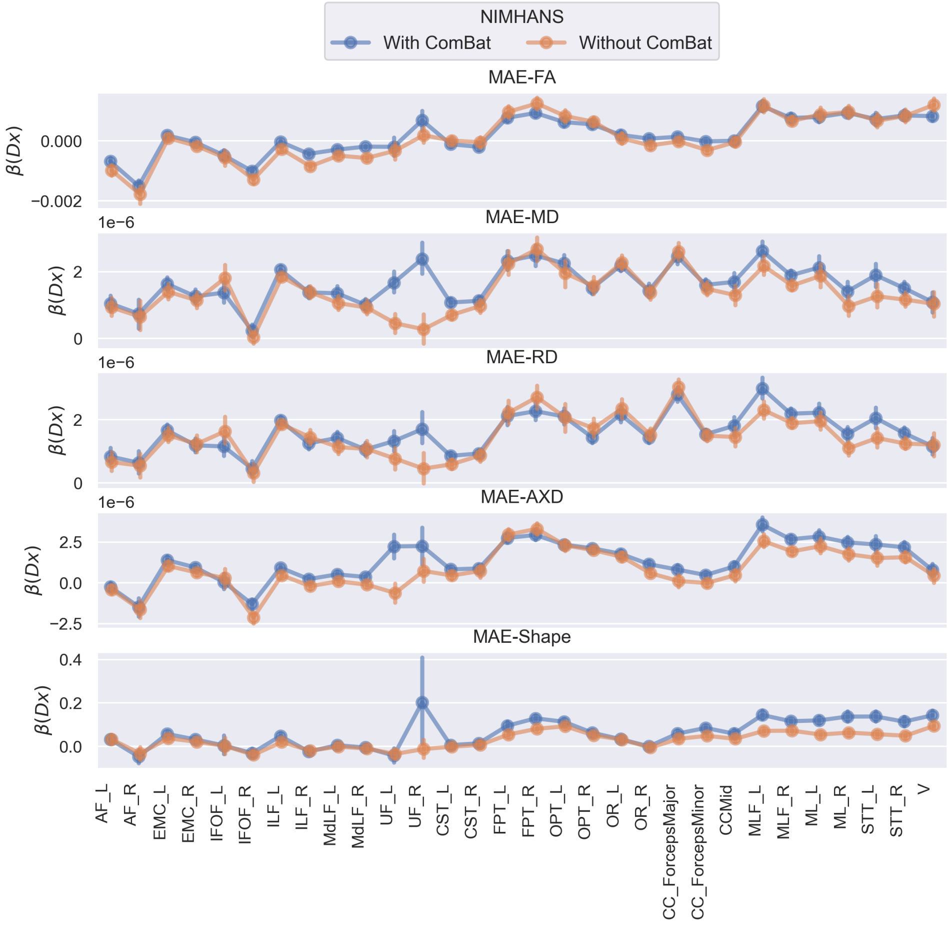

Figure 14: Fixed effect coefficient of diagnosis $\beta(\mathbf{DX})$ before and after ComBat harmonization for MAE measures calculated from subjects in the NIMHANS cohort.
